# The Transcriptional Regulator Ume6 is a Major Driver of Early Gene Expression during Gametogenesis

**DOI:** 10.1101/2023.03.28.534652

**Authors:** Anthony Harris, Elçin Ünal

## Abstract

The process of gametogenesis is orchestrated by a dynamic program of gene expression, where a vital subset constitutes the early meiotic genes (EMGs). In budding yeast, the transcription factor Ume6 represses EMG expression during mitosis. However, during mitosis to meiosis transition, EMGs are activated in response to the meiotic regulator Ime1 through its interaction with Ume6. While it is known that binding of Ime1 to Ume6 promotes EMG expression, the mechanism of EMG activation remains elusive. Two competing models have been proposed whereby Ime1 either forms a coactivator complex with Ume6 or promotes Ume6 degradation. Here, we resolve this controversy. First, we identify the set of genes that are directly regulated by Ume6, including *UME6* itself. While Ume6 levels increase in response to Ime1, Ume6 degradation occurs much later in meiosis. Importantly, we found that depletion of Ume6 shortly before meiotic entry is detrimental to EMG activation and gamete formation, whereas tethering of Ume6 to a heterologous activation domain is sufficient to trigger EMG expression and produce viable gametes in the absence of Ime1. We conclude that Ime1 and Ume6 function as a coactivator complex. While Ume6 is indispensable for EMG expression, Ime1 primarily serves as a transactivator for Ume6.

## INTRODUCTION

Gametogenesis culminates in the formation of reproductive cells via a series of highly coordinated processes driven by a dynamic and tightly controlled gene expression program. One key process in gametogenesis is meiosis, a specialized form of cell division that involves recombination between homologous chromosomes and reduction of chromosome number by half. Faithful execution of meiosis is crucial, as most human miscarriages and congenital birth defects arise from meiotic errors (Hassold and Hunt 2001; Nagaoka et al. 2012). Moreover, inappropriate activation of meiotic genes has been implicated in a range of cancer types, underscoring the significance of proper meiotic gene regulation (Lingg et al. 2022; McFarlane and Wakeman 2017; Hanahan and Weinberg 2011; Feichtinger and McFarlane 2019). Therefore, understanding the mechanisms that regulate gene expression and meiotic execution during gametogenesis is of utmost importance.

In the budding yeast *Saccharomyces cerevisiae*, gametogenesis is characterized by the activation of a series of temporally distinct gene expression clusters. The first cluster, known as the early meiotic genes (EMGs), contains evolutionarily conserved meiosis-specific genes required for DNA replication, recombination, and synapsis that ensure proper segregation of chromosomes into gametes. The coordinated expression of EMGs during gametogenesis is achieved through the common upstream regulatory sequence 1 (URS1) motif found in their promoters, which is recognized by the transcription factor (TF) Ume6 (Park et al. 1992). Ume6 interacts with three other factors – Sin3, Rpd3, and Ime1 – to toggle the expression of EMGs on or off in different developmental contexts (Bowdish and Mitchell 1993; Park et al. 1992; Washburn and Esposito 2001). This is achieved through three distinct regions in Ume6: the DNA-binding domain, the Sin3-Rpd3 histone deacetylase binding domain, and the Ime1 binding domain. Ume6 coordinates the expression of EMGs by recruiting these factors to EMG promoters.

During mitotic growth, EMGs are repressed by a complex made of Ume6 and Sin3-Rpd3. Ume6-dependent targeting of Sin3-Rpd3 to EMG promoters creates a repressive chromatin state, partly through the deacetylation of histone H4 lysine 5 (Rundlett et al. 1998; Strich et al. 1989; Vidal et al. 1991; Wang et al. 1990). In parallel, the *IME1* promoter is strongly repressed by nutritional cues, and any Ime1 protein produced is kept outside the nucleus in a cyclin-CDK and TOR-dependent manner (Colomina et al. 2003, 1999; van Werven and Amon 2011). These conditions produce cells that cannot enter meiosis and ensure separation between mitotic and meiotic events.

In respirationally-competent diploid cells, the *IME1* promoter is derepressed in response to nitrogen and glucose starvation (reviewed in van Werven and Amon 2011). Once translated, Ime1 is phosphorylated by the kinases Rim11 and Rim15 to promote its nuclear localization and interaction with Ume6 (Malathi et al. 1999, 1997; Pnueli et al. 2004; Vidan and Mitchell 1997). The *IME1* promoter itself contains a URS1 motif, which allows Ime1 to regulate its own expression (Moretto et al. 2018). Therefore, the exchange of Sin3-Rpd3 for Ime1 is a key driving force in the stimulation of EMGs and the entry of cells into the meiotic program.

The functional analysis of *UME6* has largely relied on the characterization of a null mutant, *ume6Δ*, which manifests pleiotropic phenotypes and gene expression patterns resulting from constitutive loss of EMG repression (Park et al. 1992; Strich et al. 1994; Bowdish et al. 1995; Williams et al. 2002). Therefore, the subset of genes that are directly regulated by Ume6 remains unclear. Furthermore, utilization of this null mutant has made it difficult to assess the meiosis-specific functions of *UME6* and to understand how the interaction between Ume6 and Ime1 influences EMG expression. Consequently, two distinct models have been proposed to explain the impact of Ime1 on Ume6 during meiosis. The first model suggests that Ime1, which possesses an activation domain, binds to Ume6 and transforms the complex from a transcriptional repressor into an activator, thereby resulting in EMG expression (Figure 1, top panel; Rubin-Bejerano et al. 1996; Smith et al. 1993; Washburn and Esposito 2001; Bowdish et al. 1995; Raithatha et al. 2021). The second model posits that binding of Ime1 to Ume6 serves as a signal that leads to the subsequent degradation of Ume6, thereby releasing EMG repression (Figure 1, bottom panel; Mallory et al. 2007; Law et al. 2014; Mallory et al. 2012).

**Figure 1.**
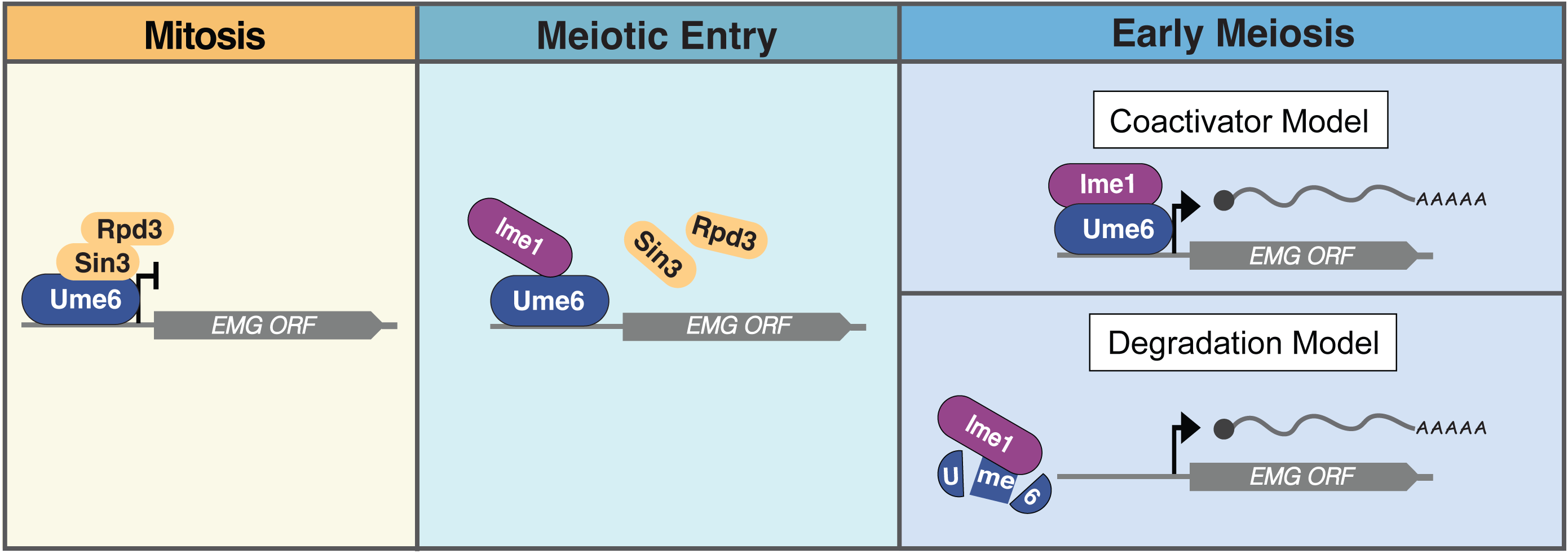
Two models of early meiotic gene (EMG) expression through Ume6 and Ime1 interaction. During vegetative growth, Ume6 associates with the Sin3-Rpd3 complex ensuring quiescence of the EMGs. The decision to enter meiosis requires exposure to nutrient and mating-type specific cues that help drive many events including: (1) dissociation of Sin3-Rpd3 from Ume6 and (2) expression of the Ime1 TF. Once expressed, Ime1 associates with Ume6. This association is critical to initiating meiotic initiation through EMG expression. However, how Ime1 binding influences Ume6 to promote EMG expression remains unclear. Two models have been presented to explain how Ime1 binding to Ume6 stimulates EMG expression. (Blue – Top) In the “Coactivator model”, Ime1 serves as a transactivator, and its binding converts Ume6 to a coactivator complex. (Blue – Bottom) In the “Degradation model”, Ime1 acts as a signal for Ume6 degradation, and its binding displaces Ume6 from EMG promoters.

By using two different meiotic synchronization methods, here we describe a thorough mechanistic characterization of Ume6’s role in meiotic gene expression. We surprisingly find that Ume6 is upregulated early in meiosis, downstream of Ime1, and is degraded only after prophase I, downstream of the transcriptional regulator Ndt80. Furthermore, by using an inducible protein depletion approach, we identify 144 genes that become derepressed upon acute removal of Ume6 in mitosis, thereby revealing its direct transcriptional targets. The expression of the same gene set is hindered when Ume6 is depleted during mitosis to meiosis transition. Thus, we provide conclusive evidence that Ume6 plays a critical role in EMG expression and gamete production, consistent with the co-activator model. This is in contrast with the role of Ume6 in mitosis, where our data confirm that it acts primarily as part of a repressive complex. Finally, by using a nanobody-based trap, we found that tethering of a heterologous transactivation domain to Ume6 is sufficient to induce EMGs and gamete production in the absence of Ime1, demonstrating that Ume6 is the primary determinant of EMG targeting. Altogether, our findings highlight Ume6 as an essential meiotic transcription factor, working in concert with Ime1, rather than a mitotic repressor that is simply an antagonist of meiotic gene expression.

## RESULTS

### Inducible depletion of Ume6 prevents the pleiotropic phenotypes associated with constitutive loss of *UME6* function

In mitotically dividing cells, Ume6 acts as part of a repressive complex leading to the silencing of EMGs (Strich et al. 1994; Williams et al. 2002). Attempts to understand Ume6’s role during mitosis have revealed hundreds of genes involved in both meiotic and metabolic functions (Park et al. 1992; Strich et al. 1994; Bowdish et al. 1995; Williams et al. 2002). However, these studies primarily relied on the use of a null mutant, *ume6Δ*, which has prolonged exposure to meiosis-specific machinery during the mitotic cell cycle, rendering it extremely sick (Figure S1A) and possibly leading to indirect effects in gene expression.

To overcome the limitations exerted by constitutive loss of *UME6* function, we utilized the auxin inducible degron system (AID, Nishimura et al. 2009), which enables rapid depletion of Ume6 carrying an AID tag (Ume6-AID) in response to the plant hormone auxin and the F-box receptor OsTIR1, which is induced by a ß-estradiol activatable transcription factor (Figure 2A, see material and methods for technical details). Cells carrying the *UME6-AID* allele grew similarly to wild type in the absence of auxin and ß-estradiol, suggesting that degron tagging of *UME6* at the endogenous locus does not interfere with its function (Figure S1A).

**Figure 2.**
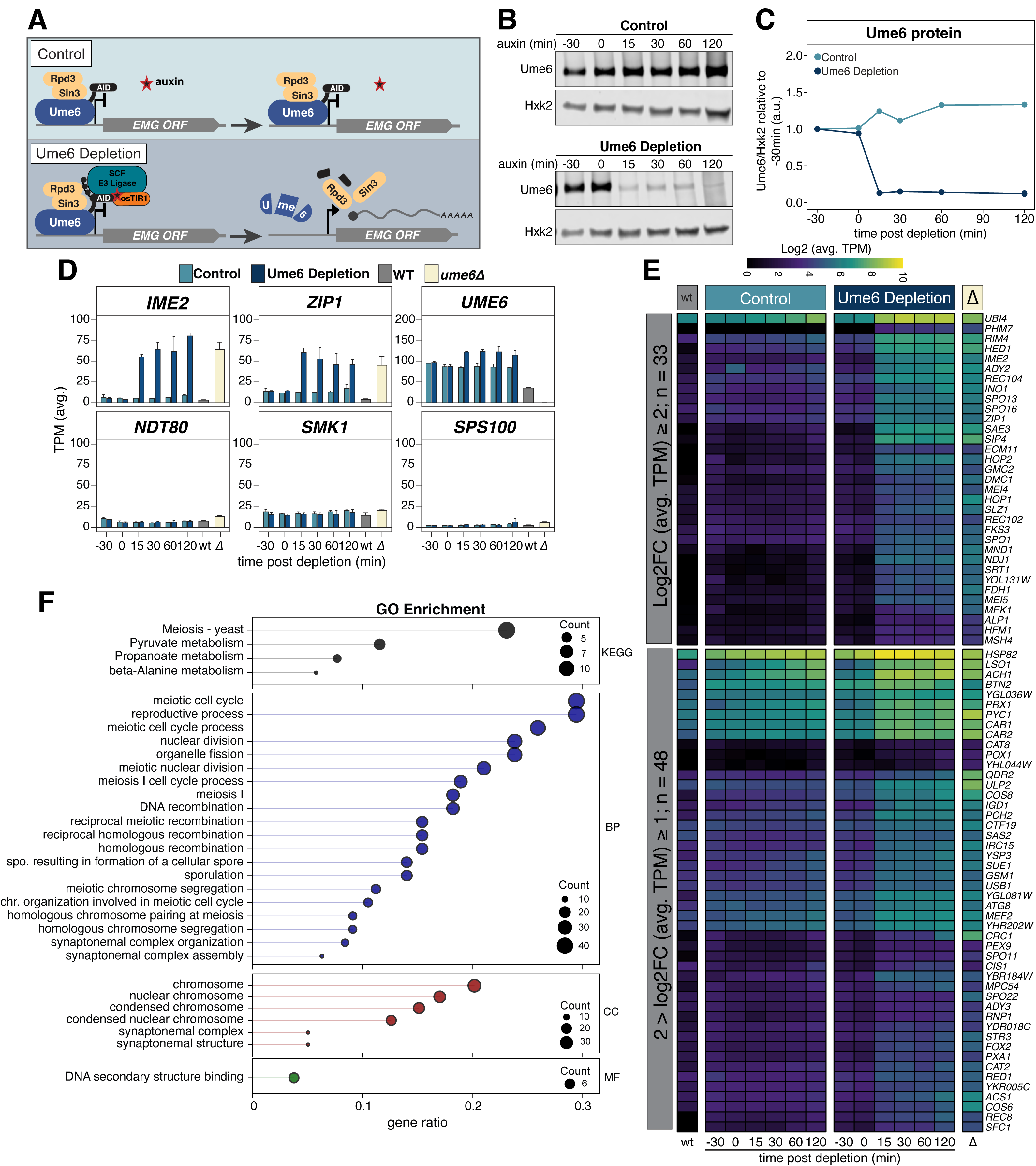
Acute depletion of Ume6 results in the derepression of EMGs in mitotically dividing cells. (**A**) Illustration of Ume6 depletion scheme using the auxin-inducible degron system. The Ume6-Sin3-Rpd3 repressive complex occupies the EMG promoters. In the absence of *osTIR* (control; light blue), the introduction of auxin doesn’t trigger Ume6 degradation and the EMG genes that it regulates remain repressed. Conversely, cells expressing *osTIR* (Ume6 depletion dark blue) in the presence of auxin recruit the E3 Ligase to the auxin-inducible degron tag associated with Ume6 for poly- ubiquitylation and subsequent degradation of Ume6-AID. Degradation of Ume6 derepresses EMGs resulting in their expression. Note that *osTIR* is under the control of *lexO* promoter, which can be induced by LexA-ER-B112 (mitotic depletion) or LexA-ER- GAL4^770-881^ (meiotic depletion) upon addition of beta-estradiol. Please refer to Materials and Methods for further details. **(B-C)** Ume6 protein levels were monitored in response to addition of auxin and beta-estradiol in the presence or absence of *osTIR* (Ume6 depletion or control, respectively). Strains possessing (UB17646) or lacking (UB18287) the *osTIR* construct or strains with wild-type *UME6* (WT; UB17716) or *ume6Δ* (*ume6Δ*; UB17718) were inoculated in YPD. Cultures were grown overnight to OD_600_ > 10 and then back diluted to OD_600_ = 0.25. Once cells reached log phase (OD_600_ = 0.5) ß-estradiol (40 nM) was added to all cultures (t_auxin_ = -30 min). Cells were allowed to continue shaking for 30 min before auxin (200 µM) was added to all cultures, initiating Ume6-AID degradation only in the *osTIR* containing strains (t_auxin_ = 0 min). Samples for protein and RNA were then collected at the designated time points. Note, cultures for wild-type *UME6* (UB17716) and *ume6Δ* (UB17718) were collected at t_auxin_ = -30 min prior to chemical treatments. **(B)** Ume6 protein levels were monitored by anti-V5 immunoblotting and using Hxk2 as a loading control. Representative blots from one of three biological replicates are shown. **(C)** Quantification of immunoblots in B. To investigate the EMG response to Ume6 degradation, RNA was extracted, and cDNA libraries were generated, sequenced, and analyzed as described in materials and methods. **(D)** Time series data for control (light blue) and Ume6 depletion (dark blue) are shown as well as for *UME6* (grey) and *ume6Δ* (ivory). The average TPMs for *IME2*, *ZIP1*, *UME6*, *NDT80*, *SMK1*, and *SPS100* are presented with standard error for three biological replicates. **(E)** A heatmap as in Figure S1E highlighting a subset of DEGs that showed the greatest response to Ume6 depletion at t = 30min. This resulted in 33 DEGs with a log2FC ≥ 2 (top) and 48 DEGs with a log2FC ≥ 1 and < 2. **(F)** GO enrichment analysis of the 144 DEGs that responded to Ume6 depletion. The gene ratio is shown on the x-axis and is the percent of genes in a given GO term out of the total 144 genes total. Point size denotes the number of genes in that GO term and color signifies category: KEGG, KEGG pathway database; BP, biological process; CC, cellular component; MF, molecular function.

To test the effectiveness of the *UME6-AID* system, we compared Ume6 levels by immunoblotting in the absence or presence of the F-box receptor OsTIR1 (from here on referred to as “control” and “Ume6 depletion”, respectively). In control cells, addition of ß- estradiol and auxin had no detectable influence on Ume6 levels (Figure 2B and 2C). In contrast, the same drug regimen resulted in the rapid depletion of Ume6 in cells carrying the F-box receptor (Figure 2B and 2C). Ume6 abundance was reduced to ∼13% of the initial levels within 15 min and remained low afterwards (Figure 2B and 2C).

To measure the transcriptomic response to Ume6 depletion, we performed RNA-seq. We initially analyzed global changes in gene expression by pairwise comparison using Spearman’s rank correlation coefficient (ρ; Figure S1B). We found that control and Ume6 depletion samples were initially very similar to one another (-30 min; ρ = 0.995). The correlation decreased, albeit slightly, following induction of Ume6 depletion (ρ = 0.978, 0.982, 0.984, and 0.984 for 15, 30, 60 and 120 min, respectively). This is perhaps not surprising given that even in the case of the *ume6Δ*, global differences in transcript levels were relatively subtle compared to wild type (ρ = 0.917). We additionally monitored sample-to-sample variation across time points using principal component analysis (PCA; Figure S1C). PC1 (58%) and PC2 (17%) accounted for 75% of the variation. Initially (-30 min), control and Ume6 depletion samples formed a distinct group, highlighting sample relatedness. After treatment with ß-estradiol and auxin, control and Ume6 depletion samples separated, with control samples only slightly shifting away from 0 min along PC1 and Ume6 depletion samples spreading along PC1 and PC2. Altogether, these transcriptome-wide comparisons indicate that gene expression patterns diverge only after induction of Ume6 depletion, thereby corroborating the temporally controlled nature of the AID system.

We next focused on the expression patterns of a subset of meiotic genes. First, we analyzed *IME2* and *ZIP1*, two well-characterized EMGs known to be repressed by Ume6 in mitosis (Figure 2D). In the control strain where Ume6 levels remained high, we observed no noticeable change in either *IME2* or *ZIP1* expression relative to wild type across all time points. However, upon Ume6 depletion, we observed a 10- and 5-fold upregulation for *IME2* and *ZIP1*, respectively (Figure 2D, 15 min). *IME2* and *ZIP1* transcripts reached similar levels to that of *ume6Δ* mutant following Ume6 depletion. Furthermore, Ume6 depletion did not affect the expression of mid meiotic (e.g., *NDT80*, *SMK1*) or late meiotic (e.g., *SPS100*) genes (Figure 2D). Together, these data suggest that the *UME6-AID* system can specifically cause derepression of EMGs in mitotic cells. Finally, we noticed reproducible upregulation of *UME6* transcripts in response to Ume6 depletion (∼30% increase between control and Ume6 depletion, Figure 2D), suggesting that Ume6 autoregulates its own expression.

### Mitotic depletion of Ume6 enables the identification of its direct targets

We next identified differentially expressed genes (DEGs) responsive to acute Ume6 depletion using two complementary approaches (see Materials and Methods for details). This analysis resulted in a composite list of 165 Ume6-responsive genes (Figure S1D; Supplemental Table 1). This list of targets was further curated using a previously published Ume6 ChIP-Seq dataset resulting in 144 Ume6-responsive genes that were also enriched for a Ume6 ChIP peak, indicating direct targets (Figure S1E, Supplemental Table 1; Tresenrider et al. 2021). To corroborate these results, we employed Multiple EM for Motif Elicitation (MEME) analysis to look for common motifs within or adjacent to the gene bodies of the 144 Ume6-responsive genes (Bailey et al. 2015; Bailey and Elkan 1994). MEME identified the core URS1 sequence (5’-GGCGGC-3) in 119 of the 144 genes (83%; Figure S1F). Further inspection of the Ume6 depletion samples revealed that all of the 144 genes were derepressed rapidly, within 15 min following auxin administration, with 58% (83/144) having a log2FC ≥ 1. Additionally, 56% (81/144) maintained a log2FC ≥ 1 at 30 min indicating a sustained expression (Figure 2E, Supplemental Table 1). In conclusion, this comparative analysis enabled the identification of a refined gene set that is directly regulated by Ume6.

Many of the Ume6 targets identified in this study are transcriptionally regulated during meiosis (Brar et al. 2012; Tresenrider et al. 2021). Brar et al. (2012) has rigorously categorized the dynamic changes in gene expression with respect to the chronology of meiotic events. Using this dataset, we determined when the Ume6 targets were expressed during meiosis. Doing so, we found that a majority reached their highest expression during meiotic entry (49/144; 34%), DNA replication (12/144; 8%), and recombination (66/144; 46%). This indicates that 88% (127/144) of our Ume6 targets are EMGs. The remaining 12% (17/144) were expressed throughout meiosis, but expression didn’t peak until mid and late meiosis suggesting additional possible layers of regulation. GO enrichment analysis for the 144 Ume6 targets was largely composed of genes involved in meiotic machinery and metabolism (Figure 2F). However, other functional classes were also revealed, including protein synthesis, trafficking, RNA processing, and cell wall maintenance. Finally, 18 genes of unknown function were present, and it is possible these genes are involved in one of the abovementioned functions that Ume6 regulates. Altogether, the comparison to the published dataset from Brar et al. confirms that the targets identified by the *UME6-AID* system represent meiotically expressed genes.

### Mitotic depletion of Ume6 derepresses meiotically-expressed LUTIs

A pervasive mechanism of gene regulation has recently been characterized in meiosis, whereby expression of a Long Undecoded Transcript Isoform (LUTI) from a distal gene promoter causes downregulation of the canonical, protein-coding transcript from the proximal promoter through the combined act of transcriptional and translational interference (Chia et al. 2017; Chen et al. 2017; Cheng et al. 2018). Among the meiotically expressed LUTIs, 72 were found to be controlled by Ume6 based on ChIP-seq (Tresenrider et al. 2021). However, a more direct interrogation of Ume6’s role in regulating these LUTIs remains unclear. Using the *UME6-AID* depletion system, we asked whether the Ume6-regulated LUTIs became derepressed in response to acute loss of *UME6* function during mitosis. Reads were aligned to a reference genome using HISAT2 and LUTI expression was monitored for all Ume6 regulated LUTIs. Of the 72 LUTIs, we identified 39 (54.2%) as being mitotically derepressed after Ume6 degradation (Supplemental Table 2). The remaining 35 failed to produce a detectable signal, possibly due to low expression and/or reduced transcript stability. Thus, our depletion system has validated a functional role for Ume6 in repressing at least 39 meiotically expressed LUTIs during mitotic growth, indicating that its activity as a transcriptional repressor can lead to both decreased and increased protein levels.

### Diametric regulation of *UME6* expression by the meiotic transcription factors Ime1 and Ndt80

The two models for Ume6-dependent control of EMG expression were postulated based on different conclusions about the levels and timing of Ume6 degradation during meiosis. These differences may have stemmed from the asynchronous nature of meiotic entry and/or the use of *UME6* null allele, which causes significant growth defects in mitosis due to EMG derepression (Strich et al. 1994; Nachman et al. 2007). To investigate the role of *UME6* in the expression of early meiotic genes (EMGs), we first followed Ume6 protein levels in a population of cells undergoing highly synchronized meiotic progression. Synchronization of meiotic progression was achieved by using an established method that utilizes a copper-inducible promoter (*pCUP1*) to control the expression of two key regulators of meiotic entry: *IME1*, which encodes an early meiotic TF, and *IME4*, which encodes an mRNA N6-adenosine methyltransferase (Berchowitz et al. 2013; Chia and van Werven 2016). We then monitored the abundance of an endogenously *3V5* tagged allele of *UME6* in these cells. Under the uninduced condition, Ume6 levels remained largely unchanged (Figure 3A and 3B, uninduced). In contrast, induction of *IME1* and *IME4* (t = 2 h) resulted in a substantial increase in Ume6 levels, up to 8-fold, which was already evident 1 h following *pCUP1* induction (t = 3 h) and remained elevated until 7 h post-induction (Figure 3A and 3B, *IME1/4* induced). Thus, Ume6 levels actually increase concurrently with EMG expression and remain elevated until around 7 h when cells transition out of prophase I.

**Figure 3.**
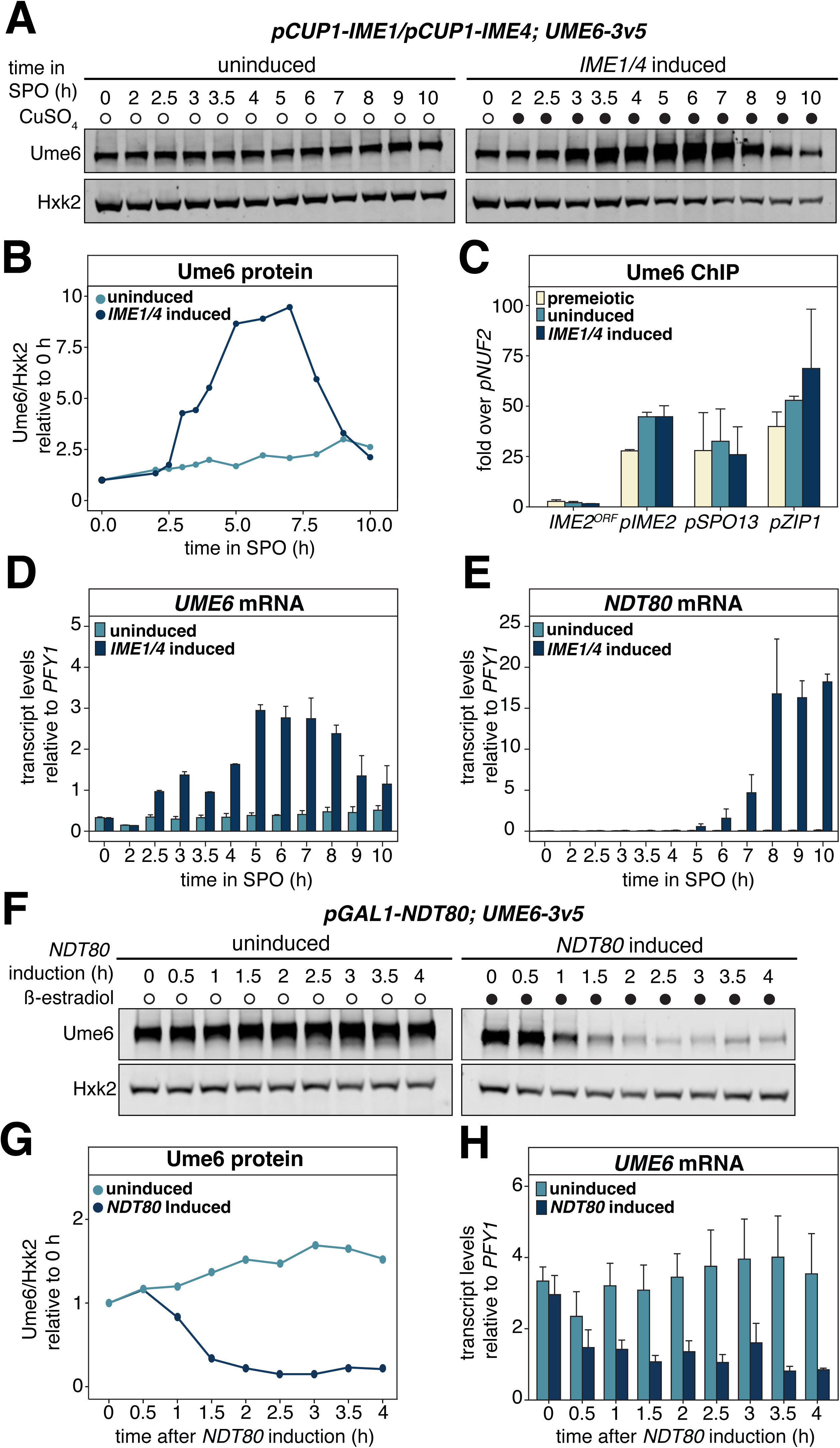
Changes in *UME6* expression in response to *IME1/4* and *NDT80* induction. (**A**) Ume6 protein abundance in response to withholding (uninduced) or inducing (induced) *IME1/4* during synchronous meiotic progression. The strain carrying *pCUP1- IME1* and *pCUP1-IME4* along with a 3v5-tagged allele of *UME6* (UB3301) was transferred to sporulation medium (SPO) at 0 h and cells were arrested at meiotic entry. After 2h, the meiotic culture was split in two. The vehicle control (water) was added to the first flask preventing meiotic entry. CuSO_4_ (50 µM) was added to the other flask to induce meiosis. Cells were collected at the indicated times for protein extraction and Ume6 levels were determined using anti-V5 immunoblotting and Hxk2 as a loading control. Representative blots from one of three biological replicates are shown. **(B)** Quantification of immunoblotting in A. **(C)** Ume6 occupancy at the *IME2*, *SPO13*, and *ZIP1* promoters, as well as the *IME2 ORF* where binding is not expected was analyzed by chromatin immunoprecipitation followed by qPCR (ChIP-qPCR) in strain UB3301. Cells were transferred to SPO and arrested at meiotic entry by withholding *IME1/4* for 2h (premeiotic). At this time, a sample of OD_600_ = 50 was collected. *IME1/4* was then either induced by addition of CuSO_4_ (50 µM; *IME1/4* induced) or withheld (uninduced). Cells were allowed to continue in SPO for 2 h after this and samples of OD_600_ = 50 for uninduced and *IME1/4* induced were collected. Mean enrichment for three biological replicates is presented with the standard error of each primer pair used. Ume6 signal at target sites was normalized over *NUF2* promoter enrichment. In addition to protein, RNA samples were collected at the indicated times to monitor expression patterns for **(D)** *UME6* and **(E)** *NDT80* in response to *IME1/4* induction. RNA was extracted from samples and transcript levels for *UME6* and *NDT80* were determined using RT-qPCR. The CT mean for two biological replicates is presented along with the range for uninduced and *IME1/4* induced samples at the specified time points. To control for technical variation, we normalized expression of *UME6* and *NDT80* relative to *PFY1*. **(F)** Ume6 protein levels in monitored in response to *NDT80* induction. The strain harboring the *pGAL1-NDT80* and *GAL4-ER* in combination with 3v5-tagged Ume6 (UB21877) was transferred to SPO. Cells were allowed to progress through meiosis for 5 h before arresting at pachytene of prophase I (t=0 h). A sample for protein and RNA was collected and cultures were split into two flasks. The first flask received the vehicle control (EtOH) to withhold *NDT80* expression (uninduced) while the other flask received ß-estradiol (1µM) to induce *NDT80* expression allowing exit from prophase (*NDT80* induced). Samples were collected at the designated time points. Ume6 levels were determined using anti-V5 immunoblotting and Hxk2 for a loading control as before. Representative blots from one of three biological replicates are shown. **(G)** Quantification of immunoblots in F. **(H)** *UME6* transcripts in the presence and absence of *NDT80* were monitored by RT-qPCR after RNA extraction. The CT mean of three biological replicates is presented along with the standard error. Technical variation was controlled for by normalization to *PFY1*.

To test whether Ume6 remained bound to EMG promoters following *IME1/4* induction, we performed chromatin immunoprecipitation followed by quantitative polymerase chain reaction (ChIP-qPCR). Ume6 enrichment was monitored at the promoters of three well-characterized EMGs, *IME2*, *SPO13*, and *ZIP1,* as well as the open reading frame (ORF) of *IME2* where Ume6 is not expected to bind. In the three EMGs analyzed, Ume6 remained bound at these promoters at levels similar to premeiotic conditions, irrespective of *IME1/IME4* induction. Thus, Ume6 is not displaced from EMG promoters during meiotic entry, suggesting that it plays a role in EMG activation during meiosis (Figure 3C).

Given that *IME1* and *IME4* are involved in transcriptional and posttranscriptional gene regulation, respectively (Shah and Clancy 1992; Hongay et al. 2006; reviewed in van Werven and Amon 2011), we reasoned that the elevated Ume6 protein levels in meiosis could be due to an increase in *UME6* mRNA abundance and hence Ume6 synthesis. To investigate this further, we analyzed *UME6* transcripts by reverse transcription and qPCR (RT-qPCR). In the absence of *IME1/4* induction, *UME6* expression was largely unchanged (Figure 3D, *IME1/4* uninduced). However, in response to *IME1/4* induction, we observed ∼7-fold increase in *UME6* expression going from pre- to post-induction. Furthermore, *UME6* expression peaked after 5 h and was maintained around this level until 8 h, consistent with the immunoblotting data (Figure 3A and 3D). Together, these findings demonstrate that *IME1/4* induces the expression of *UME6* and leads to elevated Ume6 levels during meiotic entry.

Following an initial increase, Ume6 levels began to decline after ∼7 h in SPO. This timing coincided with the expression of *NDT80* (Figure 3E), which encodes a TF necessary for exit from meiotic prophase I, initiation of meiotic divisions, and gamete maturation (Xu et al. 1995). To directly test how Ndt80 influences Ume6, we took advantage of an inducible *NDT80* system whereby *NDT80* expression is triggered by a β-estradiol-activatable TF (Benjamin et al. 2003; Carlile and Amon 2008). Cells were grown in SPO for 5 h to achieve prophase I arrest, and then β-estradiol was withheld or added to the media, thereby either preventing or allowing for *NDT80* expression and progression through the meiotic divisions, respectively. In the absence of *NDT80* induction, Ume6 levels remained unchanged (Figure 3F and 3G, uninduced). However, after 1.5 h following *NDT80* induction, Ume6 abundance was reduced to 32% of the initial levels and reached 15% after 4 h (Figure 3F and 3G, *NDT80* induced). To assess whether *NDT80* induction also influenced *UME6* transcriptionally, RNA samples were collected for RT-qPCR. Withholding *NDT80* induction resulted in *UME6* transcript levels remaining largely unchanged (Figure 3H; uninduced). In contrast, *NDT80* induction led to a ∼32% drop in *UME6* transcripts as early as 1.5 h (Figure 3H; *NDT80* induced). We conclude that Ume6 protein levels decrease in response to Ndt80, not Ime1, and this downregulation is due in part to a reduction in *UME6* transcript levels. Downregulation of *UME6* following *NDT80* expression thus restricts the timing of Ume6 removal to when meiotic cells are transitioning from early to mid-meiotic gene expression.

Cdc20, which serves as an activator for the APC E3 ligase, has been previously implicated in Ume6 degradation (Mallory et al. 2007). To shed light on Cdc20’s role in Ume6 turnover, we combined a meiotic-null allele of *CDC20*, *cdc20-mn*, with the inducible *NDT80* system *(pGAL-NDT80; GAL4-ER; pCLB2-CDC20*). However, Ume6 levels still declined in the *cdc20-mn* mutant following *NDT80* induction (Figure S2A). Thus, *CDC20* does not appear to be involved in Ume6 turnover.

Our data thus far help differentiate Ume6’s meiotic role in EMG activation through three key insights: (1) *IME1* expression results in the upregulation of *UME6* itself, leading to increased Ume6 protein levels; (2) Ume6 remains bound to the EMG promoters in the presence of Ime1; and (3) *NDT80* expression triggers events that lead to the downregulation of *UME6*, and thus reduced Ume6 protein levels, following exit from prophase I. These results are consistent with a model whereby Ime1 and Ume6 form a coactivator complex and once the early meiotic events are completed, Ume6 is downregulated in an Ndt80-dependent manner.

### Meiotic depletion of Ume6 inhibits gamete formation and prevents proper activation of EMGs

Our findings support the notion that the Ime1-Ume6 coactivator complex drives the expression of EMGs; however, it remains unclear how loss of *UME6* function, specifically during gametogenesis, impacts meiotic progression and gene expression. To address this question, we combined *UME6-AID* with the inducible *IME1/4* system, thus allowing us to rapidly deplete Ume6 shortly before meiotic entry. Our data thus far indicate that the Ume6 regulon contains at least 144 direct targets and that Ume6 is highly expressed during early meiosis, remaining bound to the EMG promoters. Thus, we predicted that depletion of Ume6 during meiosis would lead to a failure in EMG expression, disrupting gamete formation. To test the consequences of Ume6 depletion on meiosis, cells were grown as before using the inducible *IME1/4* system and were allowed to acclimate to SPO for 30 min. Ume6 depletion took place over the next 1.5 h at which point *IME1/4 was* induced. Samples for protein and RNA were collected prior to and following Ume6 depletion and *IME1/4* induction. Consistent with our previous observations, in control cells, Ume6 levels increased by 58% as early as 30 min following *IME1/4* induction and doubled by 4.5 h (Figure 4A and 4B). In contrast, cells that were depleted for Ume6 experienced a noticeable drop in Ume6 levels, down to 30.6% of starting levels at 6 h (Figure 4A and 4B; Figure S3A, please refer to materials and methods for a detailed description of the differences between mitotic and meiotic depletion strains and conditions).

**Figure 4.**
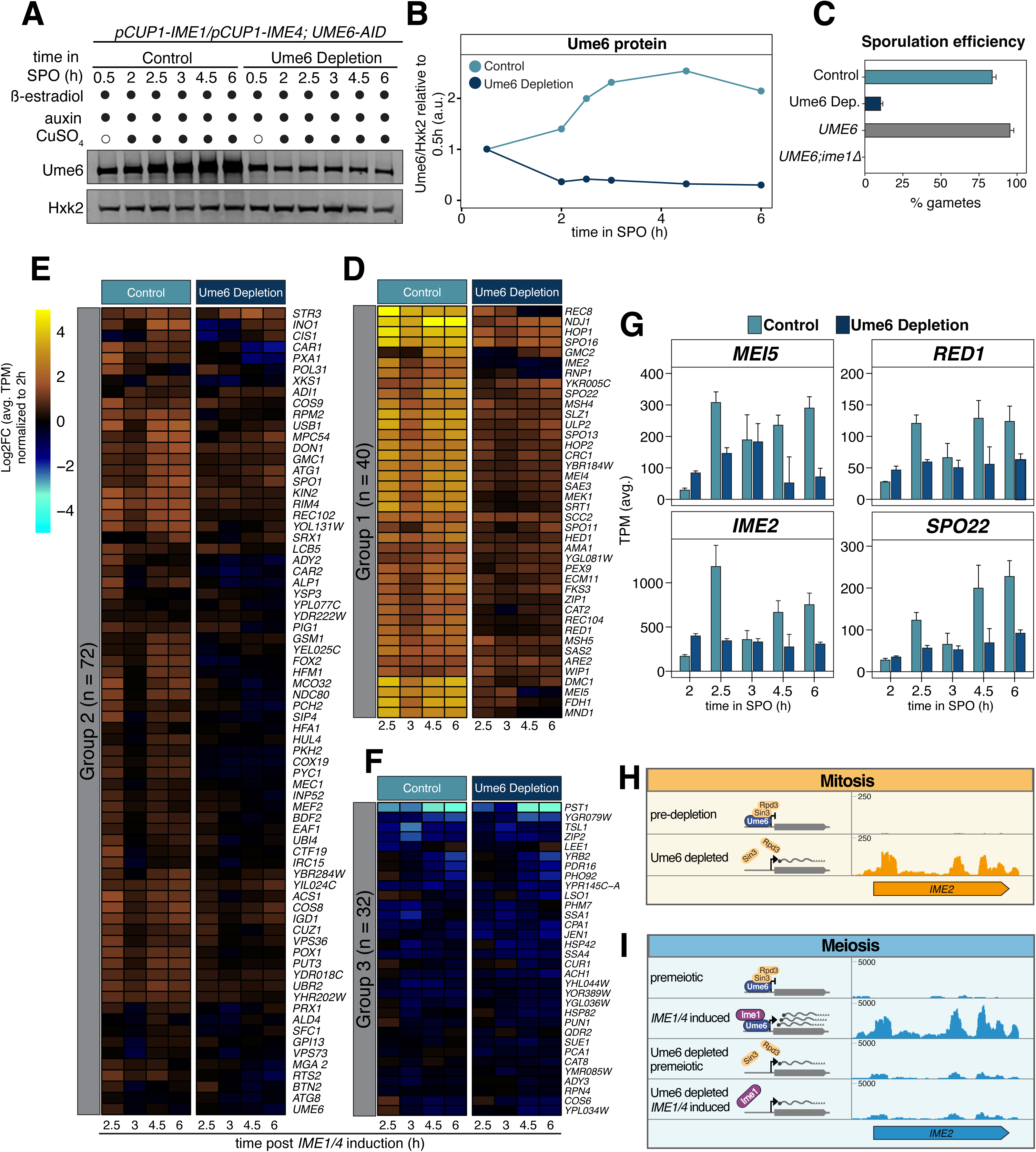
Ume6 depletion shortly before meiotic entry disrupts gamete formation and EMG expression. Cultures from control (UB25688) and Ume6 depletion (UB25092) strains were transferred to SPO at 0 h. ß-estradiol (5 nM) and auxin (200 µM) were added at 0.5 h. Then, CuSO_4_ (50µM) was added at 2 h to induce meiosis. Samples for protein and RNA were collected at designated times. **(A)** Ume6 protein abundance was analyzed using anti-V5 immunoblotting and Hxk2 as a loading control. Representative blots from one of three biological replicates are shown. **(B)** Quantification of the immunoblots in A. Sporulation efficiency data for control (UB25688) and Ume6 depletion strains (UB25092) along with two additional controls, a *pCUP1-IME1*; *pCUP1-IME4*; *UME6* (*UME6*) strain (UB19103) and *ime1Δ* (UB19105) strain. Cells were allowed to complete meiosis for 24 h before calculating sporulation efficiency. For this, 100 cells were counted and percentage of cells that formed tetrads were noted as % gametes. Data from average of three biological replicates is presented for control (light blue), Ume6 depletion (dark blue), *UME6* (grey), and *ime1Δ* (black). Error bars indicate standard error **(D-F)** The log2FC of average TPMs for three biological replicates is shown for the 144 Ume6 targets identified from mitotic cells. To evaluate EMG response to Ume6 degradation following *IME1/4* induction, TPMs were normalized to the 2 h time point just before *IME1/4* induction. Those Ume6 targets with no change in expression after *IME1/4* induction are colored black. Targets whose expression decreases are colored blue/cyan. Targets whose expression increases are colored sienna/yellow. A heatmap of the 144 Ume6 targets shown in Figure S3B was split using k-means clustering based on their Euclidean distance and application of the “elbow test” to identify an optimal k of 3 (k-means = 3). This produced three distinct groupings of genes based on their response to *IME1/4* induction: group 1 **(D)**, group 2 **(E)**, and group 3 **(F)**. **(G)** Four genes from group 1 were selected for closer inspection. This included *MEI5*, *IME2*, *RED1*, and *SPO22*, which are presented as a bar plot showing mean of TPMs including the standard error for three biological replicates. This bar plot compares control (light blue) and Ume6 depletion (dark blue) conditions at the designated times. **(H and I)** Genome browser views of mRNA tracks highlighting consequential differences between depleting Ume6 in mitosis compared to meiosis for the EMG *IME2*. **(H)** During mitosis, prior to Ume6 depletion (pre- depletion) *IME2* signal is mostly undetectable (top). Then, in response to Ume6 depletion (Ume6 depleted) *IME2* signal appears indicating a loss of repression. **(I)** Conversely, in the presence of Ume6 without Ime1 (premeiotic), signal for *IME2* is mostly undetectable, similar to mitosis, and after *IME1/4* induction (*IME1/4* induced) *IME2* becomes strongly expressed as indicated by the mapped reads. However, in the absence of Ume6, before or after *IME1/4* induction (Ume6 depleted + premeiotic and Ume6 depleted + *IME1/4* induced, respectively), *IME2* signal remains slightly stronger than premeiotic but much weaker than *IME1/4* induced. Scales for the genome browser under each condition, mitosis, and meiosis, are indicated and an illustration for Ume6’s presence and interaction with its cofactors is provided. Scales on the y-axis show relative track heights between mitosis and meiosis.

To determine the impact of Ume6 depletion on meiosis, we next analyzed the cells’ ability to produce gametes, known as spores in yeast. For comparison, a strain containing only the inducible *IME1/4* system (*pCUP1-IME1; pCUP1-IME4; UME6)* as well as an *ime1Δ* mutant (*UME6*; *ime1Δ*) where meiosis cannot occur was included (Figure 4C). *ume6*Δ cells were too sick to process for the meiotic experiments. Sporulation efficiency was 95.3% for the *pCUP1-IME1/4* strain and 0% for the *ime1Δ* mutant (Figure 4C). In the control strain where Ume6 was not depleted, sporulation efficiency was 84%, indicating that our system experiences only minor deficiencies (Figure 4C). In contrast, the Ume6 depletion strain displayed a severe reduction in sporulation efficiency (10%; Figure 4C), indicating that acute removal of Ume6 inhibits cells’ ability to complete the meiotic program.

To probe the underlying transcriptional consequences of Ume6 depletion, we performed RNA-seq and analyzed our previously generated list of 144 mitotically repressed Ume6 targets to assess whether Ume6 was necessary for their meiotic expression. We monitored the Log2FC of average TPM relative to the 2 h time point just before *IME1/4* induction and found that the majority (112/144; 78%) of the mitotically repressed Ume6 targets now showed reduced expression upon Ume6 depletion relative to the control sample (Figure 4D-F; Figure S3B-E).

To better highlight genes that are most impacted by Ume6 depletion, we applied k-means clustering, which groups genes by their Euclidian distance while minimizing variation. Using the “elbow method,” we found k = 3 to be optimal for subdividing our 144 genes in the Ume6 regulon. Group 1 contained 40 genes that were important for meiotic recombination and chromosome pairing, while group 2 had a combination of 72 meiotic and metabolic genes. Inspecting these subgroups, we found that genes in group 1 and 2 showed an average of ∼43% and ∼20% decrease in expression, respectively, in response to Ume6 depletion (Figure 4D and 4E, comparing TPM for Ume6 depletion and control at 2.5 h). Indeed, Spearman analysis at 2.5 h for group 1 and 2 showed high dissimilarity between control and Ume6 depletion (ρ = 0.435 and 0.454, respectively; Figure S3C and S3D). This reduced expression and dissimilarity between conditions persisted until 6 h. Conversely, group 3 contained several different genes involved in meiosis, metabolism, and cell wall maintenance. Expression profiles for group 3 were overall more similar between control and Ume6 depletion (∼3% increase to expression in Ume6 depletion sample compared to control at 2.5 h). Consistently, Spearman analysis showed increased sample relatedness from 2.5 h until 6 h (ρ = 0.74, 0.71, 0.86, and 0.88, at 2.5, 3, 4.5, and 6 h respectively; Figure 4F and Figure S3E). Thus, depletion of Ume6 prior to *IME1/4* induction disrupted 112 of our 144 Ume6 targets (78%) important for meiotic progression, highlighting Ume6’s importance in EMG activation during meiosis.

Focusing on four representative EMGs from group 1, *IME2*, *MEI5*, *SPO22*, and *RED1*, we found that transcriptional differences were already detectable early on, since strains retaining a functional Ume6 (control) had lower basal expression levels, consistent with Ume6 acting repressively prior to *IME1* expression (Figure 4D and 4G). Furthermore, in the control strain, gene expression spiked going from pre-*IME1/4* induction at 2 h to post- *IME1/4* induction at 2.5 h reaching 7-, 11-, 5-, and 4-fold, for *IME2*, *MEI5*, *SPO22*, and *RED1*, respectively (Figure 4G). However, depletion of Ume6 resulted in largely unchanged expression for *IME2*, *MEI5*, *SPO22*, and *RED1* (Figure 4G). Taken together, the failure to form gametes combined with reduced transcript levels of meiotic genes in response to Ume6 depletion emphasizes the critical involvement of Ume6 in the expression of EMG*s*.

These findings demonstrate Ume6’s dual role both as a repressor and an activator. By acutely depleting Ume6 under distinct developmental programs, we arrived at two very different outcomes. Mitotic depletion of Ume6 resulted in the derepression of its target genes, illustrating Ume6’s role in ensuring EMG quiescence during the mitotic gene expression program (Figure 4H). Consistently, depletion of Ume6 under nutrient-deprived conditions (i.e., in the absence of *IME1*) also led to derepression of EMGs (Figure 4I; premeiotic). However, this level of EMG derepression was not sufficient to initiate meiosis. On the other hand, depletion of Ume6 shortly before *IME1/4* induction prevented the proper activation of EMGs, thereby exemplifying Ume6‘s second role as an activator of EMGs during the meiotic program (Figure 4I). Thus, Ume6 serves as a primary determinant as to whether cells silence or induce the meiotic gene expression program depending on the cellular state and the associated cofactors.

### Tethering of Ume6^T99N^ to Ime1 using the GFP nanobody trap system rescues meiotic defects associated with *UME6^T99N^*

Previous studies have demonstrated that the meiotic kinase Rim11 phosphorylates both Ime1 and Ume6 to promote their interaction (Mitchell and Bowdish 1992; Rubin-Bejerano et al. 1996; Malathi et al. 1997). One key phosphorylation residue in Ume6 is Threonine 99 (T99). Indeed, a particular mutation at this position, T99N (Ume6^T99N^), was found to severely reduce Rim11’s ability to phosphorylate Ume6 (Bowdish et al. 1995; Malathi et al. 1997), thereby preventing binding of Ume6 to Ime1. To restore the interaction between Ime1 and Ume6^T99N^, we utilized a GFP nanobody trap approach where Ume6^T99N^ carrying a *3V5* epitope was fused to the VH16 anti-GFP nanobody (*UME6^T99N^-3V5-αGFP*; Figure 5A; Fridy et al. 2014). For controls, we included *UME6*, *UME6-3V5*, and *UME6^T99N^-3V5*. We then combined the *UME6* alleles with either *IME1* or an N-terminally GFP-tagged *IME1* at the endogenous locus (*GFP-IME1*; Moretto et al. 2018). If the interaction between Ime1 and Ume6 is sufficient to drive EMG expression, then in the *UME6^T99N^-3V5-αGFP GFP-IME1* strain, where tethering occurs, sporulation should be rescued.

**Figure 5.**
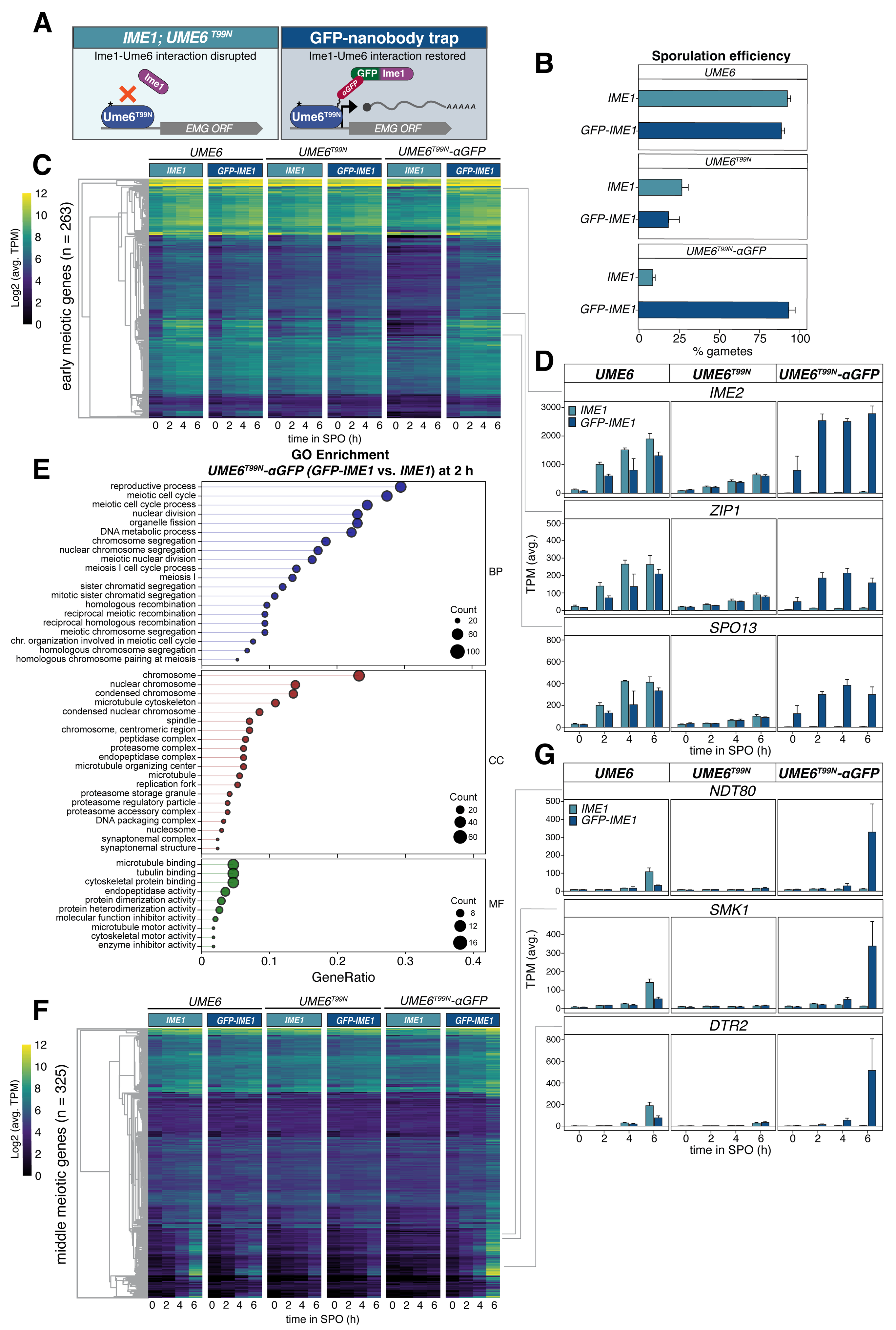
Tethering of Ime1 and Ume6 using the GFP nanobody rescues the *UME6^T99N^*meiotic defects. (**A**) Illustration of the GFP-nanobody trap approach. **(B)** Sporulation efficiency measured for strains containing either wild-type *IME1* and *UME6- 3V5* (UB26625), *UME6^T99N^-3V5* (UB26629), and *UME6^T99N^-3V5-αGFP(VH16)* (UB27313), or *GFP-IME1* and *UME6-3V5* (UB26641), *UME6^T99N^-3V5* (UB26645), and *UME6^T99N^-3V5-αGFP(VH16)* (UB27243). *3V5* is not annotated in the figure labels for simplicity. Cells were grown in presporulation media before being transferred to SPO and allowed 24 h to complete the meiotic program before sporulation efficiency was measured. As before, 100 cells were counted and percentage of cells that formed tetrads were noted as % gametes for each allele combination and the avergae of three biological replicates is presented with the standard error. **(C-G)** Strains used in Figure 5B were transferred to SPO (t = 0 h) and RNA samples were collected at the designated times. RNA samples were processed as described in Figure 3D and TPM tables were generated from three biological replicates. To examine early gene response, a set of genes identified as early expressed by Williams et al. and Brar et al. and identified as *IME1* responsive by Tresenrider et al. were termed Early Meiotic Genes and monitored in our dataset. **(C)** Heatmap representing Log2 of the mean TPMs across three biological replicates for Early Meiotic Genes. Strains harboring distinct *UME6* alleles in combination with either untagged *IME1* (light blue) or *GFP-IME1* (dark blue) are presented on top of the heatmap. (**D**) Barplot representation for mean of TPMs at a designated time point is shown for *IME2*, *ZIP1*, and SPO13. Standard error from three biological replicates is included. *UME6* alleles for each representative gene plot are shown at the top of their respective barplot. The gene represented by the barplot is shown at the top of each group and *IME1* allele is shown as either light blue (*IME1*) or dark blue (*GFP-IME1*). DESeq2 analysis between *IME1; UME6^T99N^-3V5-*αGFP (UB27313), and *GFP-IME1; UME6^T99N^-3V5-*αGFP (UB27243) at 2 h identified 316 DEGs (padj < 0.05). **(E)** GO enrichment was used on the 316 DEGs that were upregulated (log2FC > 1.5). The gene ratio is shown on the x-axis and is the percent of genes in a given GO term out of the total 316 genes total. As before, the point size corresponds to the number of genes in that GO term while color signifies category: BP, biological process; CC, cellular component; MF, molecular function **(F)** Heatmap prepared as described in Figure 5B representing a set of genes identified in Cheng et al. as responding to *NDT80* induction (Cheng et al. 2018). **(G)** Barplots as prepared as described in Figure 5C representing *NDT80*, *SMK1*, and *DTR2*.

We first examined sporulation efficiency in the strains possessing different allelic combinations of *IME1* and *UME6*. For untagged *UME6*, the sporulation efficiency was >90% when combined with either untagged or GFP-tagged *IME1* (94.3% and 97.0%, respectively; Figure S4A). *UME6-3V5* had a small drop in sporulation efficiency (92.7% and 89% for *IME1* and *GFP-IME1*, respectively; Figure 5B), suggesting the tag mildly impairs function of Ume6. However, strains with *UME6^T99N^*-*3V5* had a severe defect in sporulation efficiency (27% and 18.7% for *IME1* and *GFP-IME1*, respectively). Addition of the GFP Nanobody to *UME6^T99N^*-*3V5* (*UME6^T99N^-3V5-αGFP*) in cells containing untagged *IME1* further reduced the cell’s sporulation efficiency to 9.0%. Despite this substantial drop in sporulation efficiency, when *UME6^T99N^-3V5-αGFP* was paired with *GFP-IME1*, the sporulation efficiency was dramatically improved to 93.7% and the resulting spores were viable (Figures 5B, 6C and 6D). Thus, restoring the interaction between Ume6 and Ime1 is sufficient to complete the meiotic program and produce healthy gametes.

**Figure 6.**
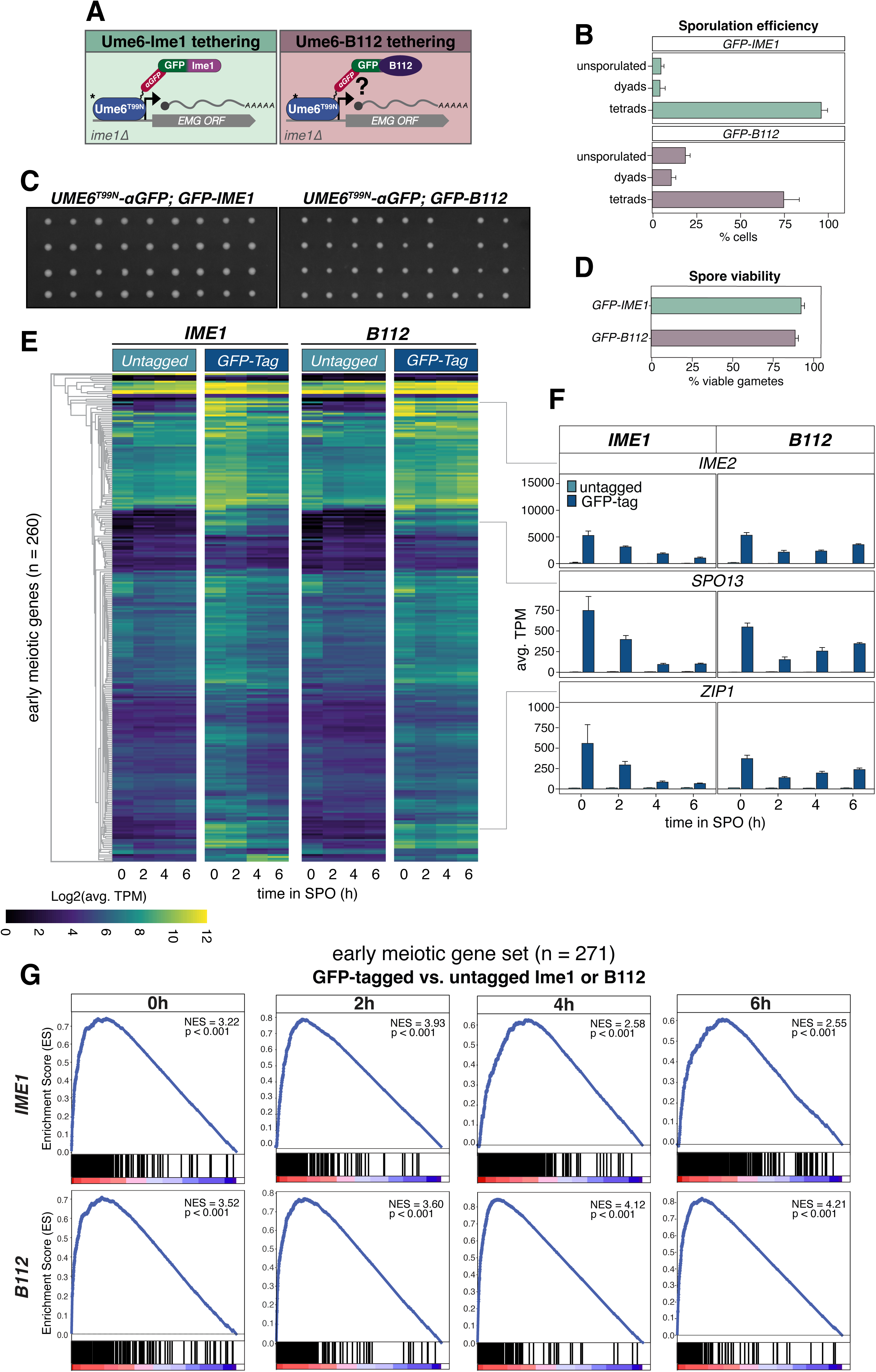
Artificial tethering of the heterologous B112 activation domain to Ume6^T99N^ is sufficient to induce meiosis and produce viable gametes. (**A**) Illustration of the GFP-Nanobody trap approach using *IME1* and the heterologous activation domain *B112* to suppress *ime1Δ* in a *UME6^T99N^-3V5-αGFP(VH16)*. Sporulation efficiency was measured for strains harboring *ime1Δ* and *UME6^T99N^-3V5-αGFP(VH16)* with either untagged *IME1* (UB32574), *GFP-IME1* (UB32572), untagged *B112* (UB33048), or *GFP-B112* (UB30295). 100 cells were counted to determine the percentage of unsporulated cells, dyads or tetrads. Data from three biological replicates along with standard error are displayed. (**C**) Strains harboring both *ime1Δ* and *UME6T99N-3V5-αGFP(VH16)* with *GFP-IME1* (UB32572) or *GFP-B112* (UB30295) were transferred to SPO and allowed to complete the meiotic program for 48 h. Spore viability was tested by digesting tetrads in zymolase 100T (1mg/ml) for 12 min before dissecting them onto nutrient rich YPD agar plates. Representative plates from the dissections are shown for *GFP-IME1*(UB32572) and *GFP-B112* (UB30295). (**D**) Quantification of spore viability. Spore viability was defined as the percent of spores that formed colonies after being transferred to nutrient rich plates out of the total 296. Note that untagged activation domains failed to produce spores and were therefore not included in the analysis. (**E** and **F**) Strains containing both *ime1Δ* and *UME6^T99N^-3V5-αGFP(VH16)* with either untagged *IME1* (UB32574), *GFP- IME1* (UB32572), untagged *B112* (UB33048), or *GFP-B112* (UB30295) were transferred to SPO (t = 0h) and RNA samples were collected at the specified times. Heatmaps for EMGs were generated as described previously and represent log2 of the mean for three biological replicates. *UME6^T99N^-3V5-αGFP(VH16)* is present in combination with *IME1* (left) or *B112* (right) either untagged (light blue) or GFP*-*tagged (dark blue) in both heatmaps. (**F**) Barplot showing mean TPMs for the indicated genes for either untagged (light blue) or GFP-tagged (dark blue) *IME1* (left column) or *B112* (right column). (**G**) Gene set enrichment analysis (GSEA) comparing untagged and sfGFP-tagged *IME1* (top) or *B112* (bottom) for “Early Meiotic Genes” at designated time points. The blue line represents enrichment for a set of genes in a given sample and the peak position denotes the degree to which that set is over- or underrepresented. The enrichment score was then normalized to account for gene set variation and is presented as normalized enrichment score (NES). Vertical black bars represent a gene and its position along the heatmap (bottom) shows how enriched that gene is in either *GFP-AD* (red, left-side) or untagged *AD* (blue, right-side).

To investigate the extent to which the nanobody-based tethering of Ume6^T99N^ to Ime1 rescues the meiotic transcriptional program, we collected RNA samples for various allelic combinations of *UME6* and *IME1* at 0, 2, 4, and 6 h. First, we inspected 263 genes that are associated with driving early meiotic events based on previous studies (Mao-Draayer et al. 1996; Pâques and Haber 1999; Williams et al. 2002; Brar et al. 2012; Tresenrider et al. 2021). 89 of these genes were present in the Ume6 direct target list (Supplemental Table 1). In *UME6-3V5* carrying either untagged or GFP-tagged *IME1*, many of these genes were upregulated after transfer to SPO (Figure 5C, compare 0 vs 2, 4, 6 h). Introduction of *UME6^T99N^*-*3V5* resulted in a moderate disruption of EMG expression (Figure S4B). Consistent with the sporulation data, *UME6^T99N^-3V5-αGFP* had a more severe defect in EMG expression than *UME6^T99N^-3V5* (Figure S4B). However, tethering of Ume6^T99N^ to Ime1 restored EMG expression back to wild type (Figure S4B). This rescue was further supported by PCA, where points associated with *GFP-IME1; UME6- 3V5 or GFP-IME1; UME6^T99N^-αGFP* separated from untagged *IME1; UME6^T99N^-αGFP* along PC1 (Figure S4C).

To globally identify the functional classes of genes expressed by tethering of Ime1 to Ume6, we performed DESeq2. Comparing *IME1* to *GFP-IME1* in the *UME6^T99N^-3V5- αGFP* background, we identified 316 DEGs (padj < 0.05; log2FC > 1.5; 2h in SPO; Figure S4G). Of the 316 DEGs identified by DESeq2, 137 were present in the EMG list and 70 were present in the Ume6 direct target list. GO enrichment revealed a number of early meiotic terms such as homologous recombination and SC formation, indicating that the tethering strategy restored early meiotic functions (Figure 5E). Additionally, inspecting tethering results for our mitotic Ume6 target list showed a similar rescue in expression (Figure S4E and S4F). Finally, key EMGs including *IME2*, *ZIP1* and *SPO13* displayed a strong rescue in their expression when Ume6^T99N^ was tethered to Ime1 (Figure 5D). We note that at 0 h, use of the GFP nanobody trap resulted in unusually high expression for many EMGs (Figure 5C and 5D). This is likely due to the high affinity between GFP and the αGFP antibody, which can bypass post-translational regulations that control Ime1- Ume6 interaction and nuclear localization, thereby resulting in earlier meiotic initiation. Regardless, these data further corroborate the significance of Ime1-Ume6 interaction in establishing the meiotic program.

We also checked the magnitude and timing of *NDT80* expression along with its targets (Figure 5F and 5G). In the *UME6-3V5* control strain, *NDT80* expression remained low from 0 to 4 h at which point *NDT80* expression increased ∼6.5 fold (Figure 5G; from t = 4h to 6h). Conversely, *NDT80* transcripts were largely unchanged in strains with *UME6^T99N^-3V5* or *UME6^T99N^-3V5-αGFP*. However, tethering of Ume6^T99N^ to Ime1 led to upregulation of *NDT80* (Figure 5G, ∼11.3-fold increase from t = 4 to 6 h in the *UME6^T99N^- 3V5-αGFP; GFP-IME1* strain). Expression of *NDT80* is indicative of chromosome segregation and gamete maturation and several genes have been identified as upregulated during these events (Winter 2012; Cheng et al. 2018). Many of the Ndt80 target genes responded to formation of the Ime1-Ume6 complex, or lack thereof (Figure 5F and 5G). Indeed, cells possessing *UME6^T99N^-3V5* or *UME6^T99N^-3V5-αGFP* failed to activate these genes or did so at a reduced level (Figure 5F and Figure S4D). In contrast, tethering of Ume6^T99N^ to Ime1 resulted in the timely activation of Ndt80 targets (Figure 5F and 5G). Altogether, these findings further emphasize the importance of Ime1-Ume6 interaction while also demonstrating that bringing Ime1 in proximity of Ume6 is sufficient to drive meiotic initiation and gamete production.

### Tethering of a heterologous activation domain to Ume6^T99N^ restores EMG expression, gamete formation, and viability

Since its initial discovery, Ime1 has been regarded as the master transcription factor in the activation of EMGs (Kassir et al. 1988). Strains lacking *IME1* (*ime1Δ*) fail to initiate meiosis and genetic screens have identified several mutations in *IME1* that disrupt meiotic initiation (Smith et al. 1993). Furthermore, mutations like *UME6^T99N^* or depletion of Ume6, which block Ime1’s ability to dock with Ume6 and localize to EMG promoter*s*, also result in meiotic failure (this study; Mitchell and Bowdish 1992). Thus, Ime1 is an essential factor in launching the meiotic transcriptional program.

On the other hand, the modularity of TFs has long been appreciated (Hahn and Young 2011). In fact, Ime1 itself can be broken into three distinct subdomains: an activation domain (AD), a nutrient-responsive domain, and a Ume6 interaction domain (Smith et al. 1993). Here, we found that tethering of Ime1 to Ume6^T99N^ is sufficient to initiate the meiotic program. We reasoned that this may occur because: (1) Ume6 needs to associate with an AD in order to function as a coactivator or (2) Ime1 has additional functions besides serving as a transactivator, which are restored upon recruitment to Ume6. To distinguish between these possibilities, we employed a heterologous AD from *E. coli*, known as B112 (Ottoz et al. 2014), and kept it either untagged or fused to GFP (Figure 6A). As controls, we used Ime1 or GFP-Ime1 (Figure 6A). Each transgene was integrated at the *HIS3* locus and was tested for its ability to suppress *ime1Δ* in the *UME6^T99N^-3V5-αGFP* background. All constructs were placed under the control of the *IME1* promoter to maintain physiological regulation and transgene expression was confirmed by immunoblotting (Figure S5A). Meiotic initiation through tethering of a heterologous AD to Ume6 would suggest that Ume6 is the primary determinant of EMG activation through its association with a transactivator. Conversely, failure to initiate meiosis would indicate a unique role for Ime1 in conducting the meiotic program.

To test whether B112 could rescue the defects associated with *ime1Δ*, we first measured sporulation efficiency in the *UME6^T99N^-3V5-αGFP* background. As expected, in strains carrying an untagged *B112* or *IME1* allele, no spores were formed (Figure S5B). *GFP- IME1* strain mostly produced tetrads (92%) and a few dyads (4%). Interestingly, *GFP- B112* also produced several tetrads (71.7%) and some dyads (10.3%) (Figure 6B). We next examined gamete viability for the strains that underwent sporulation. We found that 92.6% of the gametes from *GFP-IME1* were viable (Figure 6C and 6D). Notably, the *GFP- B112* strain also had high gamete viability of 93.2%, though colonies were marginally smaller (Figure 6C and 6D). These results demonstrate that tethering of a heterologous AD to Ume6 is sufficient to induce meiosis and generate viable gametes.

Next, we performed RNA-seq to gain insights into the underlying transcriptional response. First, we analyzed global differences between samples using Spearman’s rank correlation. We find that differences between strains carrying untagged *IME1* or *B112* transgenes were minimal (Figure S5C). However, comparison between untagged and GFP-fused transgenes showed a stark difference (Figure S5C). Interestingly, comparison of *GFP-IME1* to *GFP-B112* revealed high correlation at earlier time points (ρ = 0.960, 0.912 at 0 and 2 h, respectively), but divergence at later time points (ρ = 0.757, and 0.848 at 4 and 6 h, respectively). We observed similar patterns using PCA (Figure S5D). These results indicate that the gene expression profiles of *GFP-IME1* and *GFP-B112* start similarly but diverge from one another later in meiosis.

Next, we focused on the EMGs previously shown to respond to *IME1* induction (Tresenrider et al. 2021) and visualized them on a heatmap (Figure 6E). *GFP-IME1* and *GFP-B112* resulted in higher EMG expression compared to their untagged counterparts. However, while EMG expression in *GFP-IME1* began to decrease at 4 and 6 h, EMGs remain elevated in *GFP-B112*. Looking more closely at *IME2*, *SPO13*, and *ZIP1*, we observed a pattern where these transcripts in *GFP-IME1* and *GFP-B112* were at their highest at 0 h (Figure 6F). In the *GFP-IME1* strain, *IME2*, *SPO13*, and *ZIP1* transcript levels were reduced by nearly half every two hours. Conversely, in the *GFP-B112* strain, *IME2*, *SPO13*, and *ZIP1* transcript levels increased subtly after 2 h in SPO (Figure 6F). Furthermore, the Ume6 mitotic targets identified in this study were expressed in both *GFP-IME1* and *GFP-B112* (Figure S5E and S5F). To identify when peak expression of EMGs occurred in *GFP-IME1* and *GFP-B112*, we applied gene set enrichment analysis (GSEA; GSEA Subramanian et al. 2005; Mootha et al. 2003). GSEA revealed that EMG enrichment was highest at 2 h for *GFP-IME1* (Normalized Enrichment Score (NES) = 3.93; Figure 6G). Conversely, in *GFP-B112*, EMGs were most enriched at 6h (NES = 4.21). Thus, when both Ime1 and B112 are recruited to Ume6 through artificial tethering, cells are able to trigger EMG expression, albeit with different dynamics.

To understand transcription differences between *GFP-IME1* and *GFP-B112* at t = 6 h that may cause these discrepancies in EMG expression we performed DESeq2. DESeq2 identified 543 DEGs enriched in *GFP-IME1* compared to *GFP-B112*, several of which were MMGs including *NDT80* (padj < 0.05; log2FC > 1.5; Figure S6A). Consistent with this, GO enrichment terms were largely involved in ascospore wall development, a process controlled by *NDT80* (Figure S6B). Thus, the prolonged expression of EMGs in *GFP-B112* may relate to a delay in meiotic progression. The downregulation of EMGs and exit from meiotic prophase is largely dependent on activation of *NDT80* and its targets (Xu et al. 1995; Brar et al. 2012; Okaz et al. 2012; Chia et al. 2021). To determine whether *GFP-B112* also delays *NDT80* expression, we monitored *NDT80* expression in our dataset along with many of its downstream targets (Cheng et al. 2018). We first visualized *NDT80* and its targets by heatmap (Figure 7A). During early time points (t = 0-2 h) *NDT80* expression was low in both *GFP-B112* and *GFP-IME1* (Figure 7B). *NDT80* transcript level increased in *GFP-IME1* by ∼5.7 fold going from 2 to 4 h, whereas *NDT80* levels in *GFP- B112* only began increasing going from 4 to 6 h (∼4.5 fold). To determine whether the delay in *NDT80* expression extended to *NDT80* targets in the *GFP-B112* strain, we analyzed these genes in our dataset (*NDT80* target list was obtained from Cheng et al 2018). Using GSEA, we observed that the highest enrichment of *NDT80* targets occurred around 4 and 6 h for *GFP-IME1* (NES = 3.74 and 3.75, respectively; Figure 7C). Conversely, enrichment of *NDT80* targets in *GFP-B112* did not occur until 6 h (NES = 3.16) and was not as strong compared to *GFP-IME1* (Figure 7C). Taken together, our results indicate that *GFP-IME1* and *GFP-B112* are able to initiate the meiotic program and produce viable gametes in the *UME6^T99N^-3V5-αGFP* background. However, while *GFP-IME1* appears to achieve this in a timely manner, *GFP-B112* appears to have an extended meiotic prophase and subsequent delay in *NDT80* expression.

**Figure 7.**
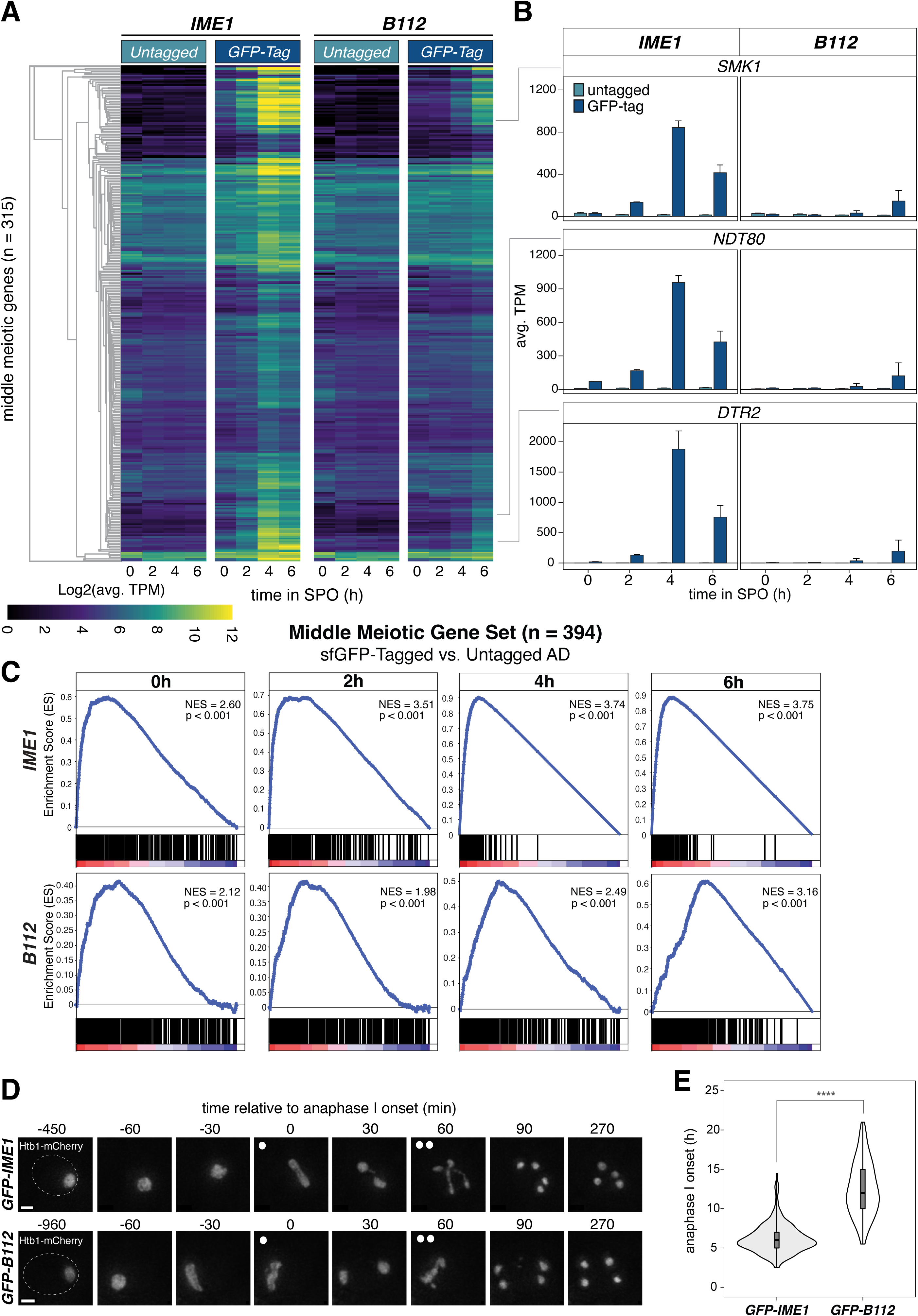
Middle meiotic gene expression is delayed with the B112 activation domain. (**A and B**) Cells were prepared as in Figure 6. Heatmaps for Middle Meiotic Genes (A and B) were made as previously described and represent log2 of the mean for three biological replicates. *UME6^T99N^-3V5-αGFP(VH16)* was combined with either *IME1* (left) or *B112* (right) lacking (light blue) or possessing (dark blue) the GFP*-*tagged, respectively. **(B)** Barplot showing mean TPMs for the indicated genes for either untagged (light blue) or GFP-tagged (dark blue) *IME1* (left column) or *B112* (right column). **(C)** GSEA was applied as in Figure 6E. This time using the NDT80 cluster from Cheng et al. 2018 to observe enrichment of *NDT80* and its downstream targets, here called “middle meiotic genes.” **(D** and **E)** Cells were transferred to SPO after brief sonication and using the CellASIC Platform in an environmentally controlled chamber at 30°C, pictures were acquired at 30 min intervals for over 21 h. **(D)** Representative images for Z-projected cells carrying either *GFP-IME1* (UB32625) or *GFP-B112* (UB31729) at the specified times. Using HTB1-mCherry, a histone marker, we labeled chromatin to identify anaphase I onset. Anaphase I is defined as the start of chromatin bifurcating into two distinct foci. Onset of anaphases I and II are denoted by either one or two white circles. Scale bar, 2 µm. Dashed lines represent cell boundary. **(E)** Quantification of anaphase I onset in cells containing *GFP-IME1* or *GFP-B112* are presented as a violin plot containing a box plot. For *GFP-IME1*, 191 cells were counted and average time to anaphase I onset was 6.2 h. For *GFP-B112*, 162 cells were counted and average time to anaphase I onset was 12.5 h. t-test results are presented on the graph and show differences in the population t(352) = -21.8, p < 0.00001 (****) in a one-tailed test.

To further confirm a delay in meiotic progression, we used an endogenously fluorescent tagged histone H2B (*HTB1-mCherry*) and performed time-lapse microscopy. We observed a delay in the onset of anaphase I in the *GFP-B112* strain (Figure 7D and 7E). While *GFP-IME1* cells took ∼6.2 h (n = 191, SD = 1.9 h) to initiate anaphase I, *GFP-B112* cells took 12.5 h (n = 162, SD = 3.4 h) to reach anaphase I (p < 0.00001; one-tailed T-test). This slowdown of meiotic progression is consistent with the delayed activation of the *NDT80* regulon in *GFP-B112*.

Taken altogether, these data support a model where Ime1’s association with Ume6 generates a coactivator complex to induce EMG*s*. Removal or disruption of the interaction between Ime1 and Ume6 hinders meiotic entry. Furthermore, EMGs can be activated when a heterologous AD is tethered to Ume6 indicating that generic transcriptional activators are able to initiate the meiotic program and even produce viable gametes when targeted to the correct genomic locations. This suggests that Ime1 serves chiefly as transactivator for Ume6 and that Ime1 has been evolutionarily tuned to allow timely expression of EMGs and execution of the meiotic program.

## DISCUSSION

The proper execution of gametogenesis relies on the timely expression of early meiotic genes (EMGs). Our study demonstrates that the transcription factor Ume6 is essential for gametogenesis. Depletion of Ume6 prior to meiotic entry disrupts EMG expression and gamete formation, highlighting its direct involvement in orchestrating the meiotic transcriptional program (Figure 4C). Notably, our study shows that the meiotic regulator Ime1 primarily functions as a transactivator within the Ume6-Ime1 complex since a heterologous activation domain from bacteria can largely substitute for Ime1 (Figure 6 and 7). Lastly, after completing early events and exiting from meiotic prophase, a second key meiotic regulator, Ndt80, facilitates the downregulation of *UME6* (Figure 3F-H). Our findings place Ume6 at the center of EMG regulation and demonstrate its essentiality in the proper execution of the meiotic transcriptional program.

### Ume6 controls its own expression through a URS1 motif

Ume6 levels remain constant in the absence of *IME1* expression. However, induction of *IME1* and *IME4* leads to a dramatic increase in *UME6* transcript levels, resulting in an upsurge in Ume6 protein abundance (Figure 3A and 3B). These findings indicate that Ume6 is not degraded in response to *IME1* expression, as previously postulated (Mallory et al. 2007; Law et al. 2014; Mallory et al. 2012), but is rather upregulated. Consistently, published datasets (Tresenrider et al. 2021; Chia et al. 2021) indicate a URS1 site proximal to the *UME6* transcriptional start site (TSS). The URS1 is located -147bp upstream of the *UME6* TSS, suggesting that Ume6 is regulating its own promoter. We further observed meiotic upregulation of *UME6* through tethering of Ume6 to a heterologous activation domain (Figure S4D). Taken altogether, Ume6 appears to stimulate its own expression during meiotic entry through a URS1 motif. Interestingly, *IME1* is also known to regulate its promoter through a URS1 motif (van Werven et al. 2012; Moretto et al. 2021). This indicates that cells have evolved feed-forward mechanisms that ensure both Ime1 and Ume6 are present at sufficient levels during meiotic entry. In addition to the feed-forward regulation by Ime1-Ume6, *IME1* mRNA is further stabilized by Ime4, which functions as an N6-adenosine methyltransferase (Shah and Clancy 1992; Hongay et al. 2006). However, it is currently unknown whether *UME6* mRNA is also regulated by Ime4. Future work could reveal more commonality between transcriptional and post-transcriptional control of *IME1* and *UME6*.

If Ume6 regulates its own promoter, then formation of the Ume6-Sin3-Rpd3 complex in mitotic cells would be expected to repress *UME6* expression during vegetative growth. Ume6 is expressed during vegetative growth, albeit at low levels (Figure 2D). Furthermore, mitotic depletion of Ume6 leads to a modest upregulation of *UME6* transcripts (Figure 2D). Thus, Ume6 appears to repress its own transcription during mitosis but to a much lesser extent than the EMGs, which are essentially quiescent in miotic cells. It has been shown that in addition to Sin3-Rpd3, Ume6 also associates with the chromatin remodeler Isw2 for full repression of EMGs (Goldmark et al. 2000; Donovan et al. 2021). Whether Ume6, Sin3-Rpd3, and Isw2 work together at the *UME6* promoter and how they simultaneously achieve silencing of the EMGs requires further investigation.

### *UME6-AID* system refines the Ume6 Regulon

The use of the AID system allowed us to deplete Ume6 in a temporally controlled manner, thereby circumventing secondary effects that arise from constitutive loss of *UME6* function (as frequently observed in *ume6Δ* mutants; Figure S1A). Importantly, combination of the depletion strategy with RNA-seq and further cross comparison to a recently published Ume6 ChIP-seq dataset enabled us to identify 144 direct transcriptional targets of Ume6 (Supplemental Table 1; Figure S1D and S1E).

Of the 144 Ume6 direct targets, 86.1% (124/144) were also present in the *ume6Δ* mutant used in this study (Figure S1G). *ume6Δ* has 1143 additional DEGs with a padj < 0.05 and log2FC > 0.6 (Supplemental Table 3). This large number of DEGs is likely due to the severe growth defect associated with *ume6Δ* and hence represents a combination of direct and indirect effects. Thus, the AID system helped to circumvent many of the pleiotropic or secondary effects associated with the *ume6Δ* mutants, while also contributing heavily to our understanding of the Ume6 regulon.

### Ume6 and Ime1 Form a Coactivator Complex to Drive EMG Expression during Gametogenesis

Our findings support the Ume6-Ime1 coactivator complex model for driving EMG expression. Besides upregulation of *UME6* following meiotic entry, four lines of evidence are consistent with this interpretation: First, depletion of Ume6 shortly before *IME1* activation disrupts EMG expression and gamete formation (Figure 4). Second, Ume6’s association with EMG promoters is unaffected by *IME1* expression as would be expected by the degradation model (Figure 3C). Third, rescuing the interaction between Ume6^T99N^ and Ime1 using a nanobody trap approach leads to an increase in Ume6 levels, rather than promoting its degradation, and enables activation of EMGs as well as meiotic execution (Figure 5). Finally, substitution of Ime1 with a heterologous activation domain from bacteria is sufficient to induce EMG expression and production of viable gametes in a Ume6-dependent manner (Figure 6). This finding also suggests that Ime1 is unlikely to possess an additional function beyond serving as a finely tuned transactivator for Ume6. Altogether, these data are consistent with a model where Ume6’s association with Ime1 forms a coactivator complex, which initiates the meiotic transcriptional program.

### The transcription factor Ndt80 downregulates Ume6 following exit from meiotic prophase

Ume6 protein levels decrease in mid rather than early meiosis (Figure 3A and 3B). This timing coincides with *NDT80* expression (Figure 3E). Ndt80 is a TF that triggers exit from meiotic prophase by inducing genes involved in meiotic divisions and gamete maturation (Xu et al. 1995; reviewed in Winter 2012). By using the inducible *NDT80* system (Carlile and Amon 2008), we show that withholding *NDT80* and arresting cells in meiotic prophase stabilizes Ume6, whereas induction of *NDT80* leads to a substantial decrease in Ume6 protein levels and downregulation of *UME6* mRNA (Figure 3F-H).

Ndt80’s involvement in *UME6* downregulation comes at a time when many early meiotic events must be terminated. It has been shown that Sin3 and Rpd3 reassociate with Ume6 later in meiosis (Pnueli et al. 2004). Furthermore, although Ime1 remains bound to Ume6 during meiosis, it has been shown that Ime1 is unable to initiate EMG expression mitotically while Sin3-Rpd3 is bound to Ume6. Reassociation of Sin3-Rpd3 with Ume6 later in meiosis could serve to downregulate Ume6 targets including EMGs and *UME6* itself (Washburn and Esposito 2001). The list of genes responsive to Ndt80 activation is quite expansive (Cheng et al. 2018). A target of Ndt80 may function to reduce or reverse the effects of Ume6 phosphorylation by Rim11 permitting Sin3-Rpd3 binding. Further investigation is required to understand the mechanism by which Ndt80 mediates *UME6* downregulation and the biological significance of such regulation.

Here we find that monitoring Ume6 turnover in a *cdc20-mn* mutant still resulted in Ume6 depletion following *NDT80* induction (Figure S2A). This is consistent with a previous report, which also found no evidence of Cdc20 involvement in Ume6 turnover during meiosis (Raithatha et al. 2021). The discrepancy is likely attributed to the use of different *CDC20* alleles. The *CLB2* promoter is strongly downregulated during meiosis thus depleting Cdc20 pools (this study and Raithatha et al. 2021). Conversely, Mallory et al. made use of a temperature-sensitive *CDC20* allele (*cdc20-ts*). The phenotype of *cdc20- ts* manifests under high temperature conditions. Exposing cells to high temperatures is known to disrupt meiotic progression in a variety of organisms. Thus, the use a *cdc20-ts* allele may have confounding effects beyond *CDC20* inactivation.

### Concluding Remarks

Our findings implicate Ume6 as a major determinant of EMG expression and successful meiotic execution. Through binding to a transcriptional activator, like Ime1, Ume6 is converted from a repressor to an activator. As part of the Ime1-Ume6 co-activator complex, both *IME1* and *UME6* appear to engage in a feed-forward mechanism by harboring a URS1-motif in their promoters. This mechanism ensures adequate protein levels and proper EMG expression through promoting their own mRNA production. Removal of Ume6 prior to *IME1* and *IME4* induction is deleterious for meiotic success and EMG expression. Furthermore, mutants that prevent proper Ime1 and Ume6 interaction also disrupt meiotic initiation. However, through reuniting Ume6 with even a heterologous activation domain, EMG expression and the meiotic program can be rescued.

This reliance on formation of a coactivator complex is functionally analogous to mammalian systems, where MEIOSIN and STRA8 form a complex to drive meiotic initiation. MEIOSIN, like Ume6, has been shown to bind to promoters of EMGs and recruit STRA8 to those sites (Ishiguro et al. 2020). Additionally, like Ime1, STRA8 has been shown to carry the activation domain necessary for EMG activation (Tedesco et al. 2009; Ishiguro et al. 2020). *Meiosin* KO strains fail to initiate meiosis even in the presence of STRA8 (Ishiguro et al. 2020). The functional similarities between STRA8-MEIOSIN and Ime1-Ume6 are striking. Therefore, our study could shed light into the transcriptional regulation of meiotic entry in more complex systems and provide a lens to investigate the associated meiotic defects.

## MATERIALS and METHODS

### Strains and Plasmids

The strains for this study, listed in **Table S4**, are derivatives of the sporulation proficient SK1 strain background (Padmore et al. 1991). The following alleles for were derived from other studies: *pCUP*-*IME1* and *pCUP*-*IME4* (Berchowitz et al. 2013), *pGAL*-*NDT80* and *GAL4*-*ER* (Benjamin et al. 2003), *HTB1-mCherry* (Matos et al. 2008), *GFP-IME1* (Moretto et al. 2018), and LexA/*lexO* (Ottoz et al. 2014).

Gene tagging or deletion was carried out using a PCR-mediate one-step integration protocol described previously (Longtine et al. 1998; Janke et al. 2004) and the PCR products generated from plasmids in **Table S5** using primers from **Table S6**.

Endogenous Ume6 was C-terminally tagged with three V5 epitopes (3V5) using plasmid pUB81 and Ume6 C-terminal tagging primers. A *UME6* degron allele was generated by C-terminally tagging endogenous Ume6 with an auxin-inducible degron (IAA7) and a 3V5 epitope from plasmid pUB763 using C-terminal tagging primers. To delete the *UME6* gene, the ORF was replaced by a HygBMX6 marker from plasmid pUB217 using *ume6Δ* primers. A plasmid containing *3V5-αGFP* for Ume6 tagging was generated as follows: a 3v5 PCR product from pUB84 was generated using 3v5 fragment primers. Along with this fragment, pUB1707 (gifted from Laura Lackner’s Lab) was subjected to *HindIII* and *Sal1* digestion at 37° for 1h. Enzymatic inactivation was then carried out at 80°C for 20min. Digested products were isolated by running them on a 1% agarose gel in 1xTBE for 25min. Fragments were then excised and transferred from the gel to a 1.5ml Eppendorf tube where they were subjected to clean up using the QIAquick Gel Extraction Kit (QIAGEN) according to protocol. Plasmid was then constructed using an NEB T4 Ligase protocol (NEB – m0202L) and transformed into competent bacteria (DH5α) for amplification. Plasmid was collected using QIAquick Plasmid Kit (QIAGEN) and named pUB2441 (*3V5-αGFP*). To C-terminally tag endogenous Ume6, a *3V5-αGFP* fragment from pUB2441 was generated using Ume6 C-terminal tagging primers.

The LexA/*lexO* system, described previously (Ottoz et al. 2014), was exploited to control *OsTIR* expression (*4xlexO-osTIR*). Additionally, to increase *OsTIR* output during meiosis, an *8xlexO-osTIR* was cloned into a *HIS3* single integration vector by Gibson Assembly (Gibson et al. 2009). Fragments were generated using pUB817, pUB99, and pUB925, with primers for *OsTIR* Fragment, *HIS* Vector, and *8x-lexO* Fragment. Fragments were then ligated according to the Gibson protocol outlined by New England BioLabs (NEB) to generate plasmid pUB2442. pUB1052 and pUB2442 were digested using *PmeI* at 37°C for 1 h. Fragments for *lexA-GAL4AD* and 8x*lexO-osTIR* were then integrated at the *TRP1* and *HIS3* locus, respectively.

Rescuing of *ime1Δ* using the heterologous activation domain (AD) B112 was achieved by constructing integration plasmids containing the full *IME1* promoter and either tagged or untagged B112. As a vector, pUB969 was amplified with pUB969 Vector Amplification primers. The fragment for *pIME1* was amplified with *pIME1* Fragment primers. Length of the *IME1* promoter was decided using Moretto et al. 2018 and ensuring both *IRT1* and *IRT2* (-2314 bp from *IME1* AUG) were included. This was done to recapitulate *IME1* transcriptional regulation and restrict AD expression to meiotic conditions. The fragment for *GFP* was amplified using *GFP* Fragment primers. A fragment for B112 with homology to *GFP* and containing the *SV40 NLS* sequence was amplified using *SV40-NLS-B112 (GFP)* Fragment primers from pUB1054. Plasmids were digested, ligated, and collected as described by the NEB protocol. Sequences were validated by PCR and sequencing and named pUB2443. The plasmid for B112 lacking *GFP* were produced using pUB2443 by first amplifying B112 using primer *SV40-NLS-B112* Fragment primers. Then, parent plasmids and fragments were digested using *XmaI* and *SacI* at 37°C for 1h before enzymes were heat inactivated at 80°C for 20min. Vector and inserts were then ligated according to the NEB protocol for the T4 ligase reaction before being named pUB2446. Single integration vectors for *IME1* were constructed in a similar way. *IME1* and *GFP- IME1* fragments were amplified from genomic DNA using primer *GFP-IME1* or *IME1* Fragment primers, respectively. Plasmid pUB2443, along with *IME1* and *GFP-IME1* fragments were digested using *XmaI* and *SacI* at 37°C for 1h before heat inactivation at 80°C for 20min. Fragments were then ligated using T4 Ligase according to the NEB protocol before being named pUB2444 (*pIME1-GFP-linker-IME1-HIS3*) or 2446 (*pIME1- IME1-HIS3*). All plasmids were sequence verified.

Ume6^T99N^ was created using a similar Cas9-based method to Sawyer et al. (2019). gRNA primers detailed in Table S6A were inserted into a centromeric plasmid (pUB1305) carrying a *URA3* marker and *pPGK1-dCas9* to generate pUB2447 and pUB2448. These plasmids were co-transformed into yeast with Ume6^T99N^ Repair Template primers to introduce the missense mutation, T99N (ACT to AAT). The plasmid was sustained on SC-ura plate for selection and successful transformants were then transferred to nutrient rich plates to lose the plasmid.

### Growth Conditions

#### Mitotic Ume6 depletion

For mitotic depletion assays, cells with wild-type *UME6* (WT), *UME6* null allele (*ume6Δ*), and *UME6-AID-3V5; lexA*-*ER-B112* strains with and without *p4xlexO-OsTIR*, were first grown in YPD (1% yeast extract, 2% peptone, 2% glucose, 22.4 mg/L uracil, and 80 mg/L tryptophan) for ∼24 h to reach saturation (OD_600_ ≥10). YPD cultures were then used to inoculate fresh YPD to OD_600_ = 0.2 and grown for ∼3 h to log phase (OD_600_ ≥0.5). During log phase, a sample for WT and *ume6Δ* was taken. Then, induction of *TIR* was initiated as follows. *UME6-AID*; *lexA*-*ER-B112* cells with and without the *p4xlexO-OsTIR* allele had *ß*-estradiol added (40 nM). Cells were incubated for 30 min before 3-indoleacetic acid (auxin) was added (200 µM). However, Ume6-AID-3v5 depletion only occurred in TIR+ strains. During the time course, samples were collected for RNA and protein extraction at -30 (ß-estradiol addition), 0 (auxin addition), 15, 30, 60, and 120 min.

#### Meiotic Synchronization

A general starvation-based method was used to sporulate cells. Briefly, cells were grown in YPD for ∼24 h shaking at 275 rpm to reach saturation (OD_600_ ≥ 10). The YPD culture was then used to inoculate BYTA (1% yeast extract, 2% bacto tryptone, 1% potassium acetate, and 50 mM potassium phthalate) to OD_600_ = 0.25 and grown for 16-18 h at 30°C to OD_600_ ≥ 5. These cells were then pelleted, washed with sterile water, and resuspended in sporulation (SPO) media (40mg Adenine Hemisulfate, 40 mg Uracil, 20 mg Histidine, 20 mg Leucine, 20 mg Tryptophan, 20g KOAc (2%) 0.02% raffinose, pH 7 in 1 L Arrowhead H2O) to a density of OD_600_ = 1.85 and shaken at 30° C at 275 rpm for the remainder of the experiment. Sporulation efficiency was always checked under a light microscope ∼24 h after shifting to SPO to determine the percentage of tetrads formed. For Figure 3A-E and 4A-G, *pCUP1-IME1* and *pCUP1-IME4* system, described previously, was used to synchronize meiotic entry as described previously (Berchowitz et al. 2013). Note that the use of *pCUP-IME1 pCUP-IME4* causes a reproducible increase in total expression many EMGs analyzed at 2.5 h before dropping at 3 h (observable in the heatmaps and individual plots). Cells appear to then equilibrate. The cause of this fluctuation is unclear but has also been observed by other researchers in the lab.

For Figure 3F-H, *NDT80* Block-Release system was used to synchronize progression into the meiotic divisions as described previously (Benjamin et al. 2003; Carlile and Amon 2008). After 5 h in SPO, *ß*-estradiol (1µM final) was added to induce *NDT80* expression. During the time course, samples were collected for RNA and protein extraction just prior to *NDT80* induction (0 h), and 0.5, 1.0, 1.5, 2.0, 2.5, 3, 3.5, and 4 h following induction.

#### Meiotic Depletion

Strains carrying both *UME6-AID-3V5; lexA-ER-GAL4^770-881^*with and without the *p8xlexO-OsTIR* allele were processed as described in “Meiotic Synchronization” with the following modifications. After 0.5 h in SPO, *ß*-estradiol (5nM final) and auxin (200 µM final) were added simultaneously. Ume6-AID-3v5 depletion occurred only in the strain carrying *p8xlexO-OsTIR*. After additional 1.5h (2h in SPO), copper (II) sulphate (50 µM final) was added to trigger *IME1* and *IME4* expression from *pCUP1* promoter to release cells into meiotic prophase. Throughout the time course, samples were collected for RNA and protein extraction: after transition to SPO (0.5 h), post Ume6-AID depletion (2 h), and post *IME1* and *IME4* induction (2.5, 3, 4.5, and 6 h). Note that the TIR+ strain used for the meiotic depletion experiments carries 8 *lexO* sites within the osTIR1 promoter and that both TIR+ (Ume6 depletion) and TIR- (control) strains contain the chimeric TF LexA-ER-GAL4^770-881^, instead of LexA-ER-B112, for triggering *osTIR1* expression in the presence of ß-estradiol. Furthermore, lower concentration of ß- estradiol (5 nM vs 40nM) was used to induce *osTIR* expression. These adjustments were necessary in order to avoid growth and sporulation defects in cells carrying *lexA-ER- GAL4^770-881^*. As a result of these modifications, the extent of Ume6 depletion was less dramatic in meiotic cells compared to mitosis. Nevertheless, we still observed significant defects in EMG expression and sporulation efficiency, indicating that meiotic cells are highly sensitive to Ume6 levels.

### Immunoblotting

For protein extraction from meiotic cultures, ∼3.7 OD_600_ of cells were collected and resuspended in 5% TCA (w/v) . For mitotic cultures, ∼ 1 OD_600_ of cells were collected. Samples were processed by centrifugation (1900 x g, 3 m, room temperature) and washed in TE50, pH 7.5 (50 mM Tris and 1 mM EDTA) and acetone before being dried overnight at room temperature. Pellets were resuspended in protein breakage buffer (TE50, 2.75 mM dithiothreitol (DTT) supplemented with 1x cOmplete EDTA-free protease inhibitor cocktail [Roche]) and disrupted using a Mini-Beadbeater-96 (BioSpec). Lysates were then mixed with 50 µL of 3xSDS loading buffer (187.5 mM Tris, pH 6.8, 6% 2- mercaptoethanol, 30% glycerol, 9% SDS, and 0.05% bromophenol blue), incubated at 95°C for 5 min to denature, and allowed to cool for at least 5 min before centrifugation at full speed for 5 min.

Proteins were separated by SDS-PAGE electrophoresis on a Bolt 4-12% Bis-Tris Plus Gel (Thermo Fisher Scientific) and then transferred onto a 0.45-µm nitrocellulose membrane in a Mini Trans-Blot Cell (Bio-Rad) containing 25 mM Tris, 192 mM glycine, and 7.5% methanol. Protein transfer was carried out using a Mini Trans-Blot Cell at a constant 180 mA (maximum, 70 V) for 3 h. Membranes were blocked at room temperature for 30 m using Odyssey Blocking Buffer (PBS; LI-COR Biosciences) before being incubated at 4°C in Odyssey Blocking Buffer (PBS) containing mouse anti-V5 antibody (RRID: AB 2556564, R960-25; Thermo Fisher Scientific) at a 1:3000 dilution for detection of 3v5 tagged alleles of Ume6. Additionally, hexokinase Hxk2 was used as a loading control and detected using a rabbit anti-Hxk2 antibody (RRID: AB 219918, 1004159; Rockland) at a 1:10,000 dilution. Membranes were incubated at 4°C for 16-18 h and primary antibody was removed. Membranes were then washed three times in 1x PBS (+0.01% Tween) shaking gently for 5 min at room temperature before being placed in the Odyssey Blocking Buffer (PBS) containing anti-mouse secondary antibody conjugated to IRDye 800CW at a 1:15,000 dilution (RRID: AB 621847, 926-32212; LI-COR Biosciences) and an anti-rabbit antibody conjugated to IRDye 680RD at a 1:15,000 dilution (RRID: AB 10956166, 926-68071; LI-COR Biosciences). Blots were washed again in PBS (+0.01% Tween-20) as before and imaged with an Odyssey CLx scanner (LI-COR Biosciences). Band intensities were quantified with the Image Studio software associated with the scanner.

### Live-cell imaging

Using CellASIC ONIX Microfluidic Platform (EMD Millipore), SPO cultures (OD_600_ = 1.85) were sonicated briefly to avoid clumping and transferred to a microfluidic Y04D plate and loaded into chambers using a pressure of 8 psi for 5 sec. Subsequently, fresh conditioned sporulation media (filter-sterilized SPO from a meiotic culture at 30°C 5 h into sporulation) was fed at a flow rate pressure of 2 psi for 24 h (King et al. 2019, 2022). The microfluidic Y04E plate was then loaded into an environmental chamber heated to 30°C mounted on a DeltaVision Elite wide-field fluorescence microscope (GE Healthcare) with a PCO Edge sCMOS camera and operated by the associated softWoRx software. Images were acquired at 60x/1.5116n oil immersion Plan Apochromat objective at 30min intervals across 21.5 h. An image stack of 4 Z positions at a 1µm step size were acquired using mCherry (10% Intensity; 25-ms exposure) and FITC (10% Intensity; 25-ms exposure) filter sets. These images were deconvolved in softWoRx software (GE Healthcare) with a 3D iterative constrained deconvolution algorithm (enhanced ratio) with 15 iterations. Once images were collected, Fiji was used to adjust brightness and contrast after images were stabilized with the Image Stabilizer plugin (Schindelin et al. 2012; Li 2008).

### RT-qPCR

For meiotic cultures, OD_600_ ∼3.7 of cells were collected. These samples were processed for total RNA first by centrifugation (2 m, 1900 g, 4° C). Supernatant was removed and cells were washed in nuclease-free water before being centrifuged again (1 min, 21000 g, 4° C). Water was removed from cell pellet and total RNA was isolated by combining acid-washed glass beads (Sigma Aldrich – G8772), 400 µL TES buffer (10 mM Tris pH 7.5, 10 mM EDTA, 0.5% SDS), and 400 µL acid phenol (0.1% w/v 8-hydroxyquinoline). The solution was shaken in a thermo mixer for 30 m at 65 °C at 1400 rpm and centrifuged (10 min, 21000 g, 4°C). Roughly 325 µL of aqueous layer was transferred to 300 µL of chloroform and centrifuged (5 min, 21000 g, room temperature). Next, 250 µL of the aqueous layer was transferred to 400 µL of 100% isopropanol (supplemented w/ 50 µL 3M NaOAc), inverted ∼10 times, and incubated for 16-18 h at 4°C. RNA was then pelleted by centrifugation (20 min, 21000 g, 4 °C) and washed in 80% EtOH. The EtOH was removed, and pellets were dried for 30-40 min before being resuspended in nuclease-free water. 5 µg of purified total RNA was then treated with DNase (TURBO DNA-free kit, Thermo Fisher (MA, USA) according to manufacturer, and 4 µL (<1 µg) of DNase treated total RNA was then reverse transcribed into cDNA (Superscript III Supermix, Thermo Fisher) according to manufacturer’s instructions. cDNA was then quantified using the SYBR green mix (Life Technologies (CA, USA)) and measured using the Applied Biosystem StepOnePlus^TM^ Real-Time PCR system (Thermofisher – 4376600). Signal for *IME2*, *NDT80*, *SPO13*, and *UME6* was measured using oligonucleotides outlined in Table S6B. Signal was then normalized to *PFY1* for meiotic cultures.

### Chromatin Immunoprecipitation

For chromatin immunoprecipitation (ChIP), meiotic culture (OD_600_ = ∼ 50) was fixed in 1.0% v/v formaldehyde for ∼20min at room temperature before quenching the reaction with 100mM glycine. Cell pellets were collected by centrifugation (3000 x g, 5min, 4°C) and washed in cold PBS. Cell pellets were then resuspended in 1 ml FA lysis buffer (50 mM Hepes pH 7.5, 150 mM NaCl, 1 mM EDTA, 1% Triton, 0.1% sodium deoxycholate) with 0.1% sodium dodecyl sulfate (SDS) and 10% w/v cOmplete^TM^ protease inhibitor pellet. Cells were broken using a mini beadbeater (BioSpec) and lysate was transferred to a 1.5 mL low adhesion Eppendorf tube and debris was cleared by centrifugation (2000 x g, 3min, 4°C). Supernatant was transferred to a new 1.5 mL low adhesion Eppendorf tube and lysate was centrifuged (20000 x g, 15min, 4°C). Supernatant was discarded, leaving a cloudy pellet behind. Pellets were resuspended in 1ml lysis buffer + 0.1% SDS + cOmplete protease inhibitor and chromatin was sheared by sonication using a Bioruptor (Diagenode (Seraing, Belgium), 8 cycles of 30sec ON/45sec OFF). Extracts were incubated overnight at 4°C in agarose beads conjugated to anti-V5 antibody (Millipore Sigma – A7345). Bead bound chromatin was then washed twice in 1ml lysis buffer, buffer 1 (250mM NaCl in lysis buffer + 0.1% SDS), and finally buffer 2 (10 mM Tris pH 8, 250 mM LiCl, 0.5% NP-40, 0.5% deoxycholate sodium, 1 mM EDTA) before reverse crosslinking was done in Tris-EDTA buffer (100 mM Tris pH 8.0, 10 mM EDTA, 1.0% v/v SDS) at 65°C overnight. After 1h of proteinase K treatment at 65°C, samples were cleaned using QiaQuick PCR cleanup (Qiagen – 28106) and enrichment of Ume6 at *IME2*, *SPO13*, and *ZIP1*, promoters as well as the *IME2 ORF* was measured by real-time PCR using SYBR green mix. Oligonucleotide sequences used for ChIP are outlined in Table S6B.

### RNA-seq

RNA samples were collected and processed as described in RT-qPCR section. To prepare mRNA-seq libraries, 10 µg of total RNA was polyA-selected and processed using the NEXTFLEX Rapid Direction RNA-seq Kit (NOVA-5138-10 and NOVA-5138-11; PerkinElmer) according to the provided manual. Quantification of resulting cDNA yields was performed using a Qubit 3 (ThermoFisher Scientific) using the high sensitivity DNA assay kit. AMPure XP beads (A63881; Beckman Coulter) were using during size selection (200-500bp) and fragment quality and quantity was analyzed using high sensitivity D1000 ScreenTapes on the Agilent 4200 TapeStation (Agilent Technologies, Inc.). Samples were sequenced through the Vincent J. Coates QB3 Genomics Sequencing Facility at the University of California, Berkeley using 100 bp single-end sequencing on an Illumina Novaseq 6000. Alignment of sequenced reads was carried out using either HISAT2 or Kallisto. For HISAT2, the protocol outlined by Pertea et al. 2016 was used with SK1 reference genome, sourced from the Saccharomyces Genome Resequencing Project at the Sanger Institute to visualize transcript isoforms (Pertea et al. 2016). For Kallisto, pseudoalignments were carried out according to a manual developed by Bray et al. to generate TPM and raw counts tables (Bray et al. 2016). Kallisto quant settings were adjusted to -b 5 -l 160 -s 20 - -single - -threads 4 based on fragment lengths determined by the Agilent 4200 TapeStation (Agilent Technologies, Inc.)

### Heatmaps and Plots

R was further used for Spearman’s correlation and with the packages pheatmap (ver. 1.0.12) and ggplot2 (ver. 3.4.0) to generate heatmaps and plots used in this manuscript, respectively (Wickham et al. 2019).

### Differential Gene Expression Analysis

Identification of differentially expressed genes (DEGs) responsive to Ume6 mitotic depletion was performed using two complementary approaches in R (R: ver. 4.1.3; RStudio: ver. 2022.07.1 Build 554). First, raw counts generated from Kallisto were exported to R. Then, using the DESeq2 package (ver. 1.34.0), differences in expression between control and Ume6 depletion samples across time (t = 0, 15, 30, 60, 120 min) were determined using an FDR of 5% (R Core Team; Love et al. 2014). To set up time series analysis, the DESeq2 “test” parameter was set to “Likelihood Ratio Test (LRT)” that, by default, uses the Wald test to generate results tables. Time series analysis between control and Ume6 depletion conditions using DESeq2 identified 177 Ume6- responsive genes (Supplemental Table 1). Depletion of Ume6 during mitosis should derepress its targets, therefore we inspected the list of 177 Ume6-responsive genes looking for sustained derepression from 15 min to 120 min (post-Ume6 depletion). We noted some genes that were largely unchanged post-Ume6 depletion (i.e. *ERO1* and *BOI1*, ∼1% and ∼3% increase, respectively, comparing Ume6 depletion to control at t = 15 min) that were counted significant by DESeq2. Thus, to control for any false positives in our list of 177, we generated a custom R script to filter out these transcripts. In brief, to avoid transcripts that displayed a response independent of Ume6 depletion, we performed pairwise analysis using DESeq2 at t = -30 min and 0 min. Those genes with differential expression at these times (padj < 0.05; abs(log2FC) > 1) were removed. Next, we looked transcripts that displayed an acute response to Ume6 depletion at 15 min (padj < 0.05; abs(log2FC) > 0.3), as would be expected of direct regulation by Ume6. Filtering of transcripts using this script reduced the list from 177 to 135.

Second, we noted that some genes previously identified in Williams et al. 2002 as being derepressed in *ume6Δ* were not present in the list of 177 (i.e. *PIG1*). Further inspection in our TPM table revealed *PIG1* did experience a ∼34% increase in expression post-Ume6 depletion (t = 15 min, comparing Ume6 depletion to control). Thus, to identify any genes missed by DESeq2, we performed additional analysis using TPM data and a custom R script. Briefly, we took the ratio between Ume6 depletion and control samples at each time point (t = 15, 30, 60, and 120 min). This was done for the TPM of all genes. Next, we took the average (avg.) of these ratios across all time points. We then looked for genes whose avg. TPM ratio between Ume6 depletion and control across time was ≥ 1.4 or ≤ 0.6. Doing so, we identified 128 genes, 98 that were previously called significant by DESeq2. The 30 additional genes were inspected before being added to the list of 135 DESeq2 targets resulting in a total list of 165. Thus, between DESeq2 and TPM analysis we identified 165 distinct genes that responded to Ume6 depletion, referred to herein as our “composite list” (Supplemental Table 1).

### ChIP Peak Curation

Using a previously published dataset from Tresenrider et al. (2021) we analyzed ChIP peak scores for our composite list of 165 Ume6 targets. We divided the average ChIP peak score (n = 3 biological replicates) for each of the 165 Ume6-responsive genes by the ChIP peak score of *IME2*, a well-characterized Ume6 target and selected those with ratios ≥ 0.5. This analysis resulted in 144 Ume6-responsive genes that were also enriched for a Ume6 ChIP peak, indicating direct targets (Figure S1E, Supplemental Table 1).

### GO Enrichment

Go enrichment was performed in R using the clusterProfiler package (Yu et al. 2012) together with the org.Sc.sgd.db (ver. 3.14.0; Carlson 2021).

### Motif Discovery

Motif enrichment analysis for Ume6 targets was performed using MEME (ver. 5.5.1 (Bailey et al. 2015). Sequences for 1,000 bp up- or downstream as well as the ORF were obtained using YeastMine (Balakrishnan et al. 2012) and exported to MEME as Fasta files for analysis with restricting the motif length’s upper limit to 15 nucleotides, but otherwise using default settings. A p < 0.05 for a motif in a given gene was considered significant. These motifs were also validated using ChIP-Seq data from Tresenrider et al. 2021.

### Gene Set Enrichment Analysis (GSEA)

Normalized counts generated from DESeq2 were compared between samples using GSEA v4.3.2 [build: 13] to assess enrichment of gene sets (Mootha et al. 2003; Subramanian et al. 2005). The “Early Meiotic Gene” set was generated by analyzing previously established data (Brar et al. 2012; Chia et al. 2021; Tresenrider et al. 2021). First, genes whose TPMs changed in response to *pCUP1-IME1 pCUP1-IME4* induction by a log2FC > 1.0 were taken from Tresenrider et al (609 genes). Next, using a list of *NDT80* targets generated in Cheng and Otto et al, we removed MMGs from this list. The remaining 518 genes were then curated using Brar et al. and Chia et al., limiting expression timing to between meiotic entry and prior to metaphase I. Finally, genes with high TPM levels during mitotic growth were also excluded. This resulted in a list of 272 early expressed meiotic genes termed “Early Meiotic Genes”. As mentioned, the second set of genes termed “Middle Meiotic Genes” were defined in Cheng and Otto et al. as a set of 394 genes responsive to *NDT80* induction (*NDT80* cluster, Cheng et al. 2018). The desktop version of GSEA was used to load in data and determine enrichment with the following modifications: “Collapse/Remap to gene symbols” was set to “No Collapse” and “Permutation Type” was set to “Gene Set”, other settings were unchanged.

### Resource availability

All reagents used in this study are available upon request from the corresponding author. Sequencing data generated in this study are available are available at NCBI GEO under the accession ID: GSM7083402 through GSM7083623

The custom code used for the analysis is available in the following code repository: https://github.com/harranth

## Acknowledgements

We thank Gloria Brar, Doug Koshland, Peter Sudmant, Kathleen Ryan, Folkert van Werven, Andrea Higdon, Amanda Su, Amy Tresenrider, Ben Styler, Cyrus Ruediger, Emily Powers, Grant King, Jingxun Chen, Jessica Leslie, Kate Morse and Tina Sing for suggestions and comments on this manuscript, Amy Tresenrider, Peter Sudmant, Christopher Mugler, Grant King, Jingxun Chen, Lizet Reyes, and Tina Sing for technical support, Jeremy Thorner and all members of the Ünal and Brar labs for valuable discussions. This work is supported by funds from the National Institutes of Health (R01 GM140005) and Astera Institute to EÜ. A.H is supported by a Chancellor’s Fellowship endowed by the University of California, Berkeley, and by an NSF-GRFP (DGE-1752814).

## Author contributions

A.H., conceptualization, data curation, formal analysis, investigation, methodology, validation, visualization, and drafting of the manuscript. E.Ü., conceptualization, visualization, supervision, project administration, funding acquisition, and drafting of the manuscript.

