## Supplemental figure legends for "The Transcriptional Regulator Ume6 is a Major Driver of Early Gene Expression during Gametogenesis"

**Supplemental Figure 1.** (Related to Figure 2) **(A)** Growth phenotype of the haploid strains *UME6* (UB17716), *UME6-AID-3V5* (UB18216), *UME6-AID-3V5; lexA-ER-B112* (UB18287), *UME6-AID-3V5; lexA-ER-B112; 4xlexO-OsTIR* (UB17646), *ume6Δ* (UB17718). Strains were serially diluted onto plates containing nutrient-rich media with agar (YPD) and allowed to grow at 30˚C for 48 h before images were acquired. Using DESeq2, counts data were analyzed for the control and Ume6 deplete time course data from Figure 2. **(B and C)** Sample-to-sample variation was analyzed using Spearman rank order correlation (ρ) with the corrplot package and the variance stabilizing transformation (VST) and plotPCA function associated with DESeq2. **(B)** Results of Spearman analysis. Red boxes are samples with low relatedness and blue boxes are samples with increased relatedness. **(C)** A PCA plot was then generated from these data using the ggplot2 package to visualize the influence of PC1 (x-axis) and PC2 (y-axis) on samples. To distinguish between samples, control (UB18287) and Ume6 deplete (UB17646) conditions were differentiated by shapes while specific time points were assigned colors. We note that replicate 3 from Ume6 depletion sample (60 min time point) is separated from the other two replicates. This is possibly due to the higher read depth in this sample. **(D)** Volcano plots were used to observe significance with which genes were derepressed (y-axis) and how this corresponded to log2FC (x-axis). Genes were either considered down- (blue) or upregulated (red) if padj < 0.05 and log2FC > 0.6. Otherwise, changes were deemed not significant (black). Dots represent a gene and are color matched to their differential expression profile. Relevant dots are provided a box denoting their gene name. **(E)** Differentially expressed genes (DEGs) between Ume6 depletion and control as well as WT and *ume6Δ* conditions were identified with the R package DESeq2 using Kallisto generated counts tables. The log2 of mean TPMs across three biological replicates are shown for the 144 DEGs identified ranging from no expression (black) to high expression (yellow). DEGs were clustered by Euclidian distance (centroid) and partitioned vertically by strain background. **(F)** The URS1 motif identified in the regions flanking or internal to the CDS of a subset of the 144 DEGs. DESeq2 was used to compare WT and *ume6Δ* (padj < 0.05; Log2FC > 0.6) resulting in identification of 1267 DEGs. (G) Venn diagram showing the 124 overlapping DEGs between Ume6 deplete (144 DEGs) and *ume6Δ* (1267 DEGs).

**Supplemental Figure 2.** (Related to Figure 3) A strain harboring the *pGAL1-NDT80* and *GAL4-ER* system was combined with the *cdc20-mn* allele (UB22674) and grown as in Figure 3F and 3G. Protein samples were collected at the designated time points and Ume6 levels were determined by anti-V5 immunoblotting using Hxk2 as a loading control.

**Supplemental Figure 3.** (Related to Figure 4) Increased levels of ß-estradiol in the presence of LexA-ER-Gal4.AD is toxic. **(A)** Growth phenotype associated various concentrations of auxin and/or ß-estradiol. Genotype of diploid strains are *UME6* (UB19103), *UME6-AID-3V5* (UB19101), *UME6-AID-3V5; lexA-ER-GAL4.AD* (UB25688), *UME6-AID-3V5; lexA-ER-GAL4.AD; 8xlexO-OsTIR* (UB25092), *ime1*∆ (UB19105), and *ume6*∆ (UB22812). Strains were serially diluted onto plates containing nutrient-rich media with agar (YPD) and allowed to grow at 30˚C for 48 h before images were acquired. **(B)** The gene set derived from mitotic depletion of Ume6 was observed by heatmap for the indicated meiotic samples. The log2 of mean TPMs across three biological replicates are shown for the 144 DEGs and normalized to t = 2 h just before *IME1*/4 induction. Expression ranges from decreased expression (cyan) to increased expression (yellow) with no expression change in black. DEGs were clustered by Euclidian distance (centroid) and partitioned vertically by strain background. **(C-E)** Heatmap for Spearman correlation between samples was generated using the corrplot package in R for group 1 (Figure S3C), group 2 (Figure S3D), and group 3 (Figure S3E). Red boxes signify poor sample relatedness while blue boxes signify high sample relatedness.

**Supplemental Figure 4.** (Related to Figure 5) (A) As in Figure 5B, sporulation results are reported for the control strain containing untagged *UME6* with either *IME1* (light blue; UB26621) or *GFP-IME1* (dark blue; UB26637). Briefly, Cells were grown in presporulation media, then transferred to SPO, and allowed 24 h to complete the meiotic program before 100 cells were counted. The average of three biological replicates is presented with the standard error. For B and D**,** Corrplot package was used in R to generate heatmaps for Spearman correlation between samples. Red boxes show samples with poor correlation and blue shows samples with high correlation. **(B)** Heatmap showing sample relatedness for the early meiotic gene set derived from Brar et al. 2012, Cheng et al. 2018, and Tresenrider et al. 2021. **(C)** Results of PCA visualized using the ggplot2 package and observing PC1 (x-axis) and PC2 (y-axis). Sample-to-sample variation by PCA was monitored for *GFP-IME1; UME6-3V5* (UB26641), *IME1; UME6$^{T99N}$-3V5*-αGFP (UB27313), and *GFP-IME1; UME6$^{T99N}$-3V5*-αGFP (UB27243) and strains were distinguished by shapes. Time points were assigned a unique color. Note that due to poor read depth, one replicate for *GFP-IME1; UME6-3V5* is isolated from the rest (4 h). **(D)** Heatmap showing sample relatedness for the middle meiotic gene set found in Cheng et al. 2018. Rescue of Ume6$^{T99N}$ using GFP Nanobody decreases variation with wild-type strains. DESeq2 results were processed using the VST and plotPCA functions associated with DESeq2. **(E)** Heatmap for the log2 of average TPM across three biological replicates of the 143 Ume6 targets. **(F)** We note that *SAE3* was dropped while constructing our heatmap due to low expression and instead is presented as a barplot here. **(G)** Volcano plot representing DESeq2 analysis between *IME1; UME6$^{T99N}$-3V5*-αGFP (UB27313), and

*GFP-IME1; UME6$^{T99N}$-3V5*-αGFP (UB27243) at 2 h that showed DEGs (padj < 0.05) either up- (red) or downregulated (blue) or showing no overall change (black).

**Supplemental Figure 5.** (Related to Figure 6) A heterologous activation domain (AD) rescues Ume6$^{T99N}$ and ime1Δ triggering meiosis using the GFP Nanobody system. **(A)** Ume6 levels were measured in *GFP-IME1* (UB32572) and *GFP-B112* (UB30295) strains by collecting time points at the designated times and performing anti-V5 immunoblotting using Hxk2 as a loading control. Values below blots were calculated by first normalizing Ume6 levels to Hxk2 in each lane, and then dividing that ratio by the initial (0 h) time point. **(B)** Sporulation efficiency measured as outlined in Figure 5B for strains with both *ime1*Δ and *UME6$^{T99N}$-3V5-αGFP(VH16)* with either untagged *IME1* (UB32574) or untagged *B112* (UB33048). 100 cells were counted and their ability to produce dyads or tetrads as well as those that remained unsporulated is presented for three biological replicates along with the standard error **(C)** Corrplot package was used in R to generate heatmaps comparing sample relatedness between *GFP*-tagged and untagged alleles of *IME1* and *B112*. Red denotes poor correlation and blue is high correlation. **(D)** DESeq2 results between untagged- and GFP-tagged ADs were processed by using VST and plotPCA functions associated with DESeq2. A PCA plot that highlights PC1 (x-axis) and PC2 (y-axis) was built using ggplot2 and is presented here. Shapes denote distinct genotypes while colors denote time points. We note that differences exist between samples at 0 h for untagged and tagged *IME1* and *B112*. This is most likely caused by the affinity GFP has for the nanobody trap, resulting in strong recruitment of the activation domain to Ume6 and earlier than normal meiotic initiation. **(E)** Heatmap showing log2 of mean TPM across three biological replicates for 143 Ume6 targets. **(F)** We note that *SAE3* again was dropped while constructing our heatmap due to low expression present and instead is presented as a barplot.

**Supplemental Figure 6.** (Related to Figure 7) **(A)** Volcano plot highlighting DEG found using DESeq2 analysis between *GFP-IME1* (UB32625) and *GFP-B112* (UB31729) at 6 h that showed 543 DEGs (padj < 0.05; log2FC > 1.5) either up- (red) or downregulated (blue) or showing no overall change (black). **(B)** GO enrichment applied to the 543 DEGs identified by DESeq2. Gene ratio is presented on the x-axis and is the percent of genes in a given GO term out of the total 543 genes total. Again, point size signifies the number of genes in that GO term while color signifies category: BP, biological process; CC, cellular component; MF, molecular function.
