## Supplementary figures and images for "The Transcriptional Regulator Ume6 is a Major Driver of Early Gene Expression during Gametogenesis"

### Figure 2 supplement

#### Harris and Ünal_Supplemental Figure 1

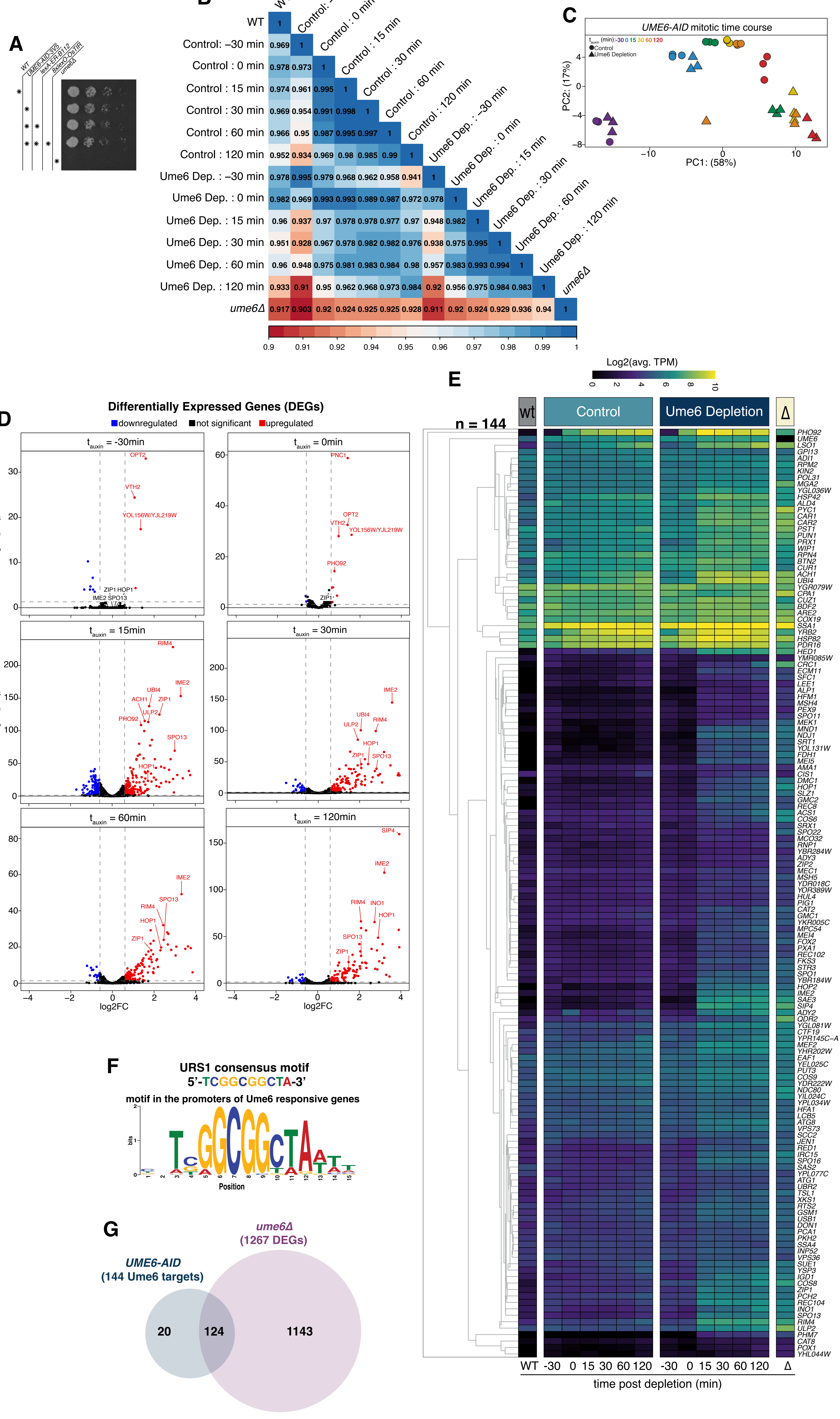

### Figure 3 supplement

#### Harris and Ünal_Supplemental Figure 2

**A**

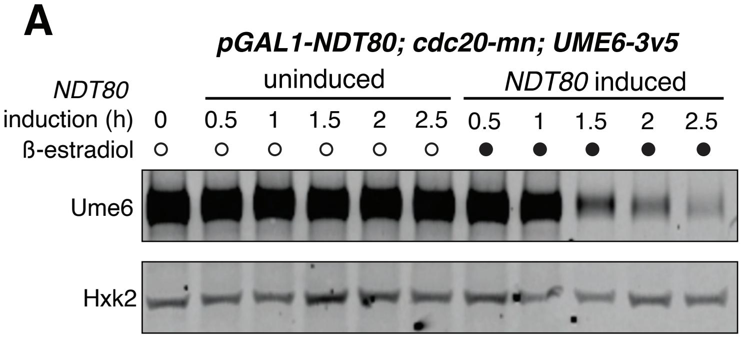

### Figure 4 supplement

#### Harris and Ünal_Supplemental Figure 3

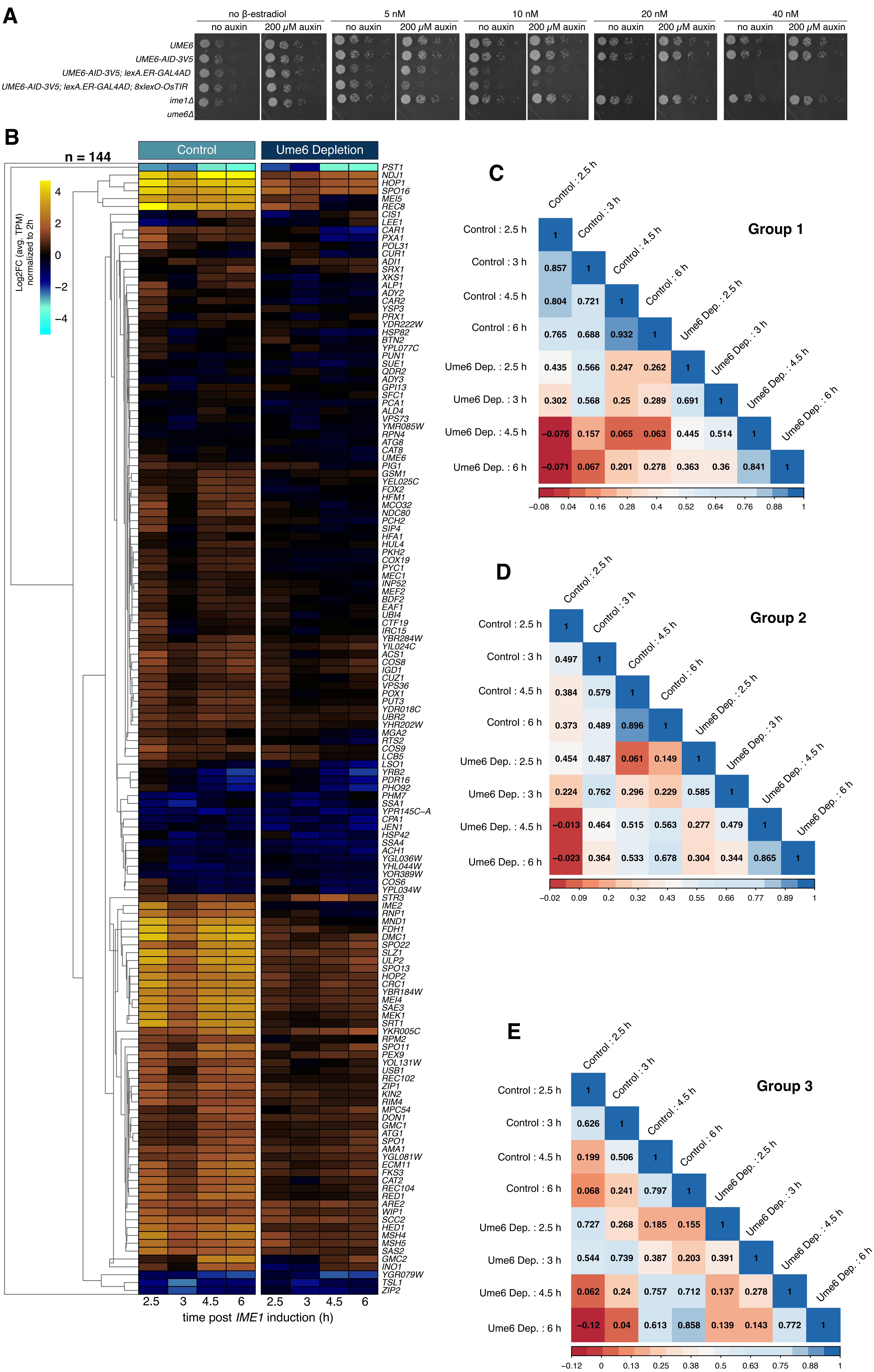

### Figure 5 supplement

#### Harris and Ünal_Supplemental Figure 4

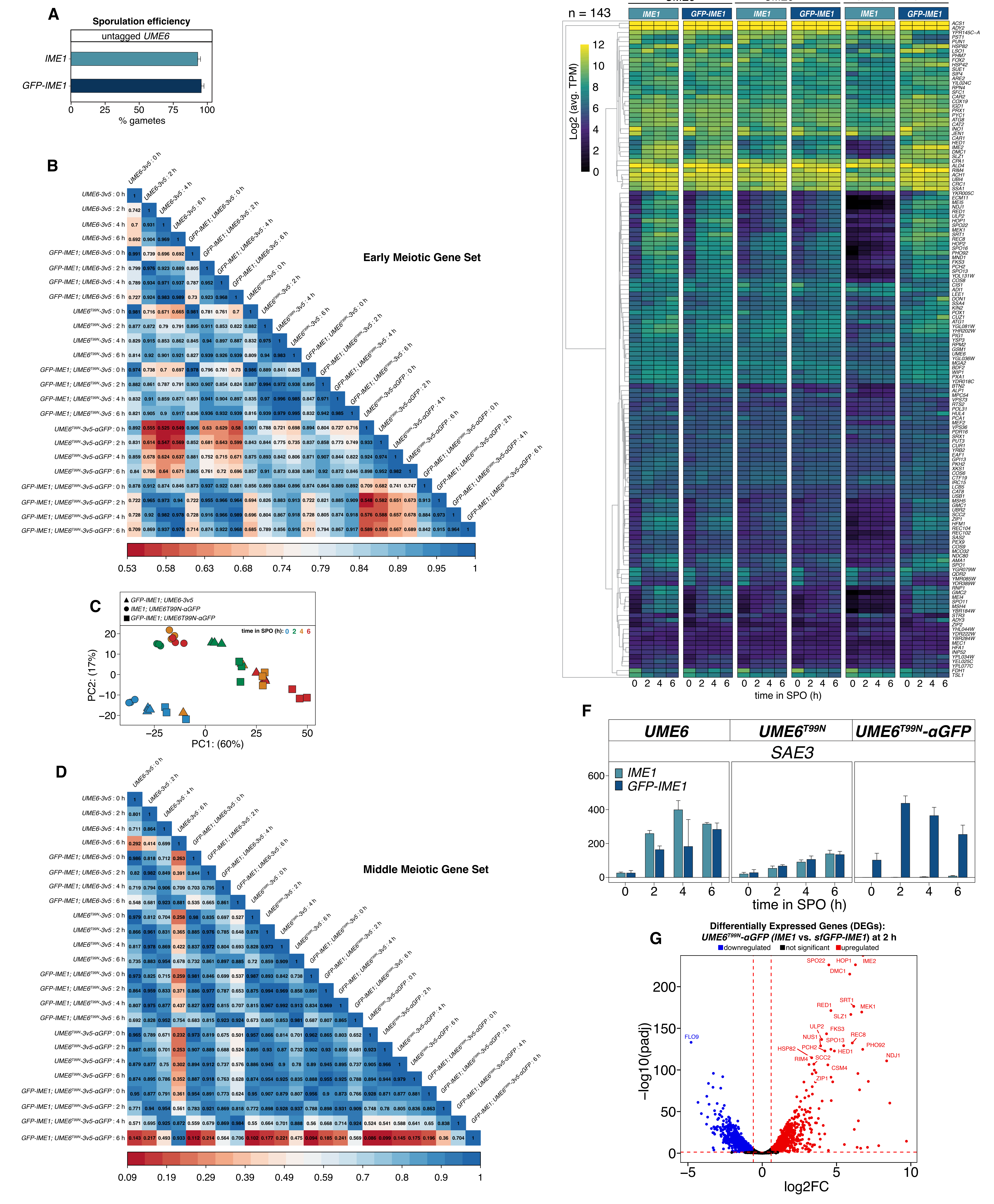

### Figure 6 supplement

Harris and Ünal_Supplemental Figure 5

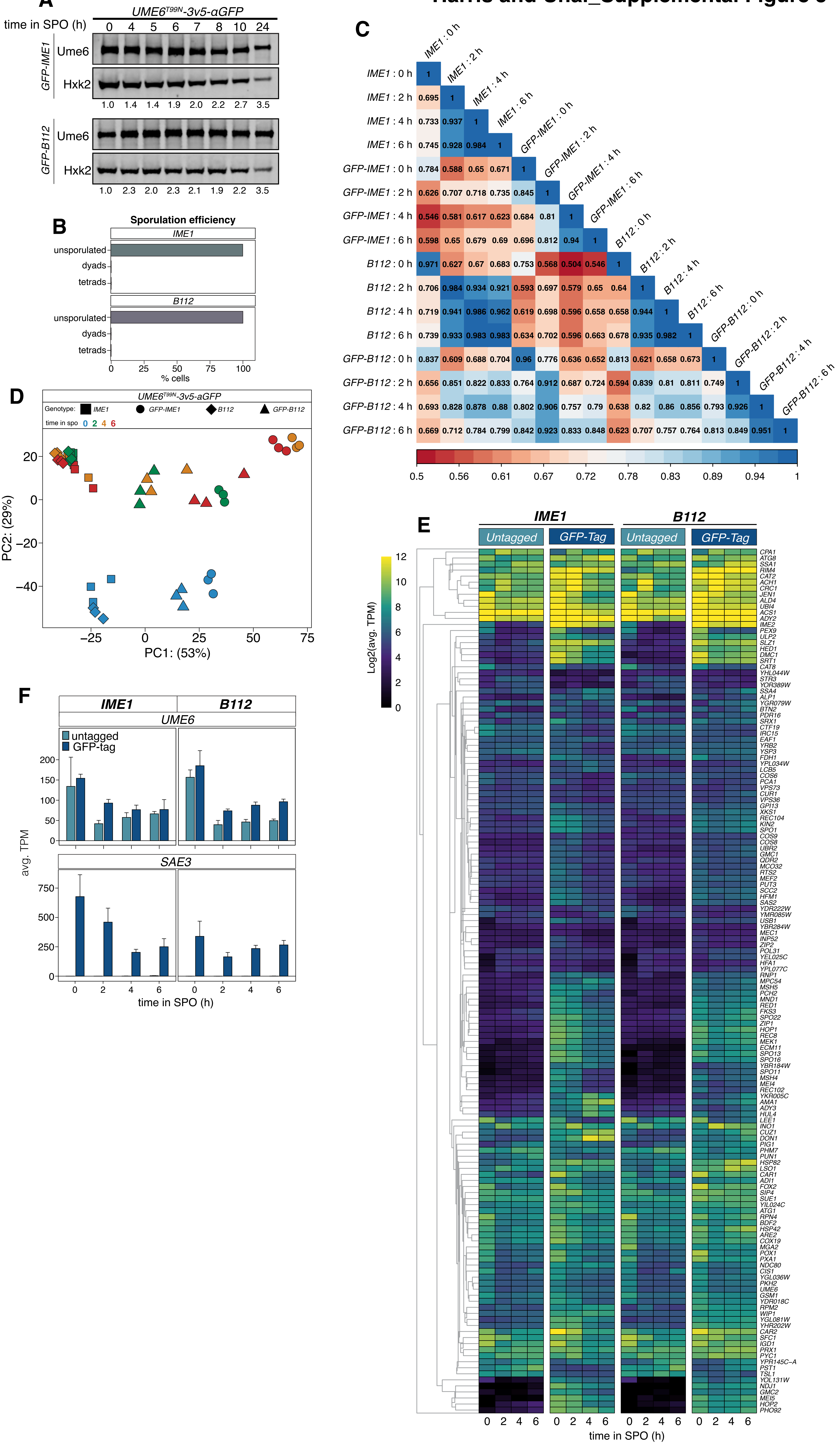

### Figure 7 supplement

#### Harris and Ünal_Supplemental Figure 6

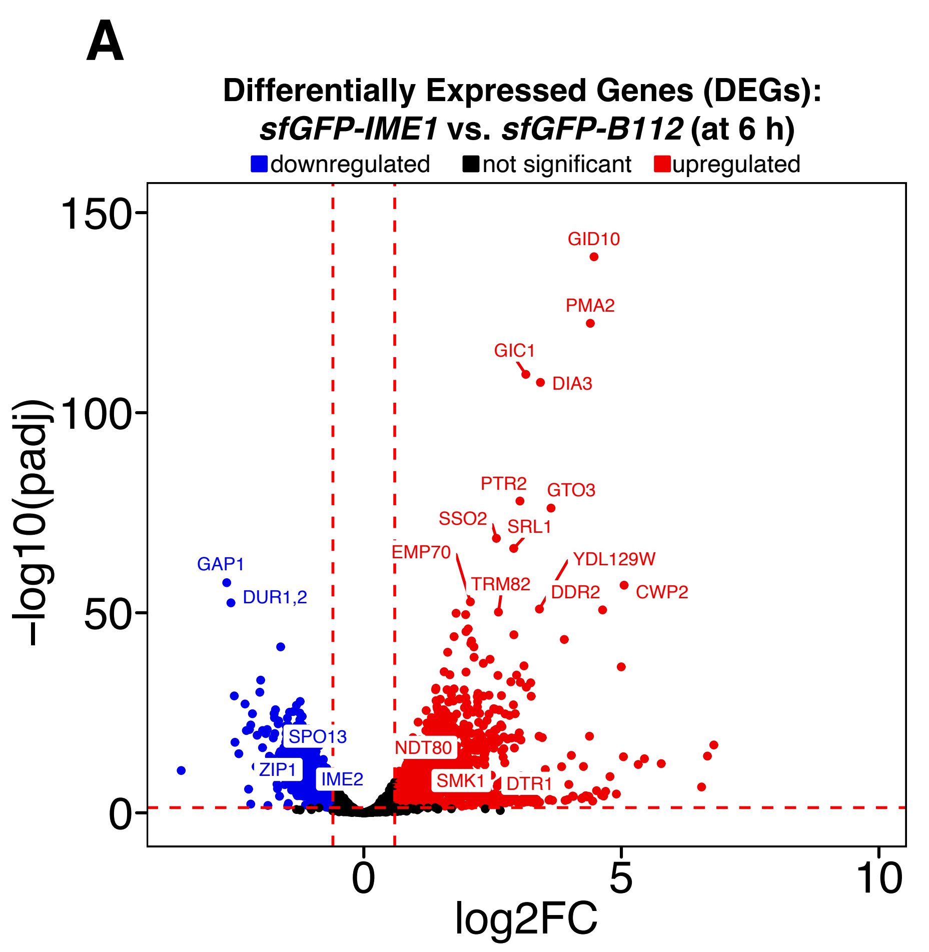


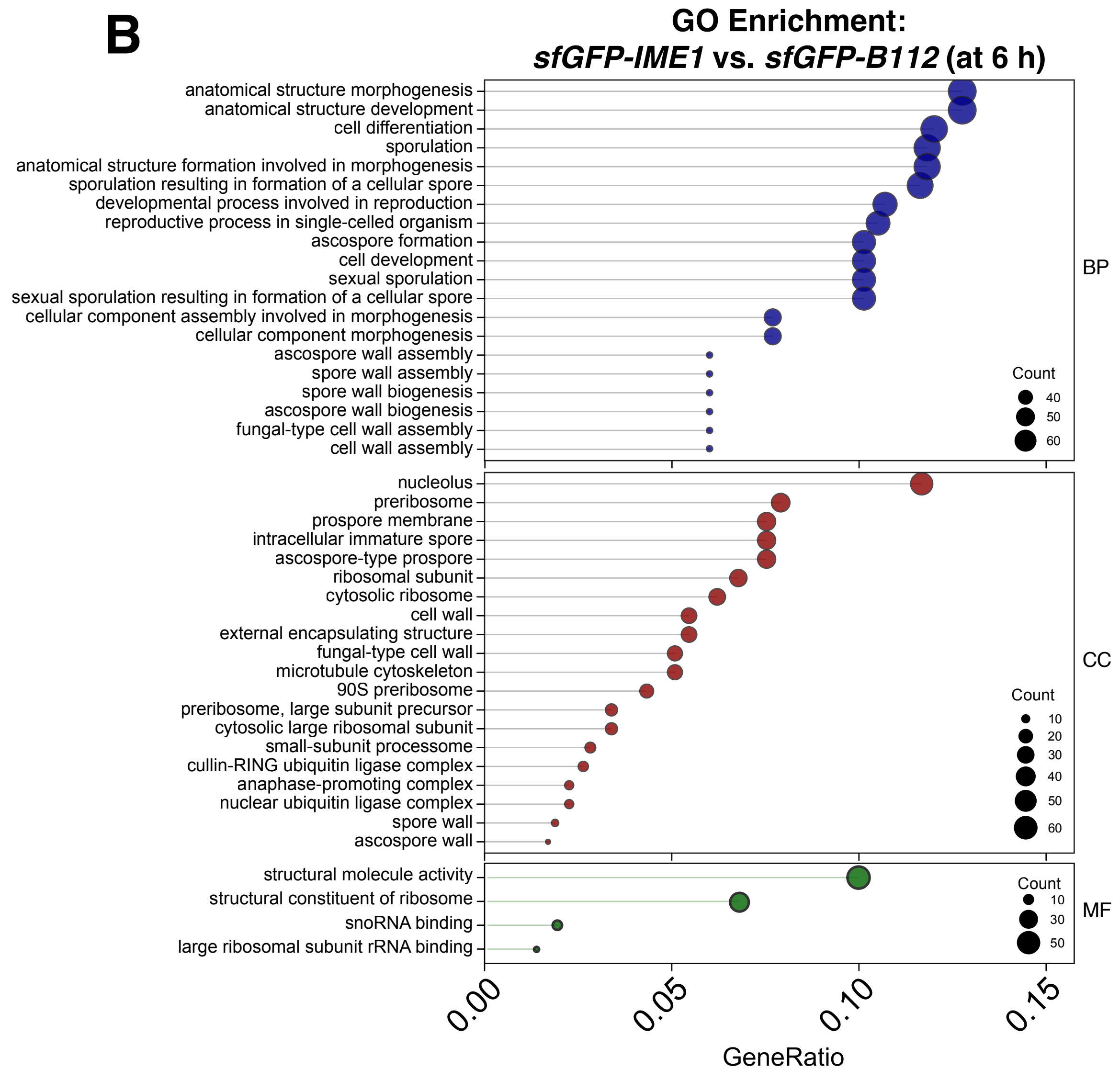
